# Identification of robust RT-qPCR reference genes for studying changes in gene expression in response to hypoxia in breast cancer cell lines

**DOI:** 10.1101/2024.08.02.606329

**Authors:** Jodie R. Malcolm, Katherine S. Bridge, Andrew N. Holding, William J. Brackenbury

**Affiliations:** Department of Biology, University of York, York, YO10 5DD, United Kingdom; York Biomedical Research Institute, University of York, York, YO10 5DD, United Kingdom; Centre for Blood Research, University of York, York, YO10 5DD, United Kingdom

## Abstract

Hypoxia is common in breast tumours and is linked to therapy resistance and advanced disease. To understand hypoxia-driven breast cancer progression, RT-qPCR quantifies transcriptional changes important for malignant development. Reference genes (RGs) are endogenous RT-qPCR controls used to normalise mRNA levels, allowing accurate assessment of transcriptional changes. However, hypoxia reprograms transcription and post-transcriptional processing of RNA such that favoured RGs including *GAPDH* or *PGK1* are unsuitable for this purpose. To address the need for robust RGs to study hypoxic breast cancer cell lines, we identified 10 RG candidates by analysing public RNA-seq data of MCF-7, T-47D, MDA-MB-231 and MDA-MB-468 cells cultured in normoxia or hypoxia. RT-qPCR determined RG candidate levels in normoxic breast cancer cells, removing *TBP* and *EPAS1* from downstream analysis due to insufficient transcript abundance. Assessing primer efficiency further removed *ACTB, CCSER2* and *GUSB* from consideration. Following culture in normoxia, or acute or chronic hypoxia, we ascertained robust non-variable RGs using RefFinder. Here we present *RPLP1* and *RPL27 as* optimal RGs for breast cancer cell lines cultured in normoxia or hypoxia. Our result enables accurate evaluation of gene expression in hypoxic breast cancer cell lines and provides an essential resource for assessing hypoxia’s impact in breast cancer progression.

**Graphical Abstract:** 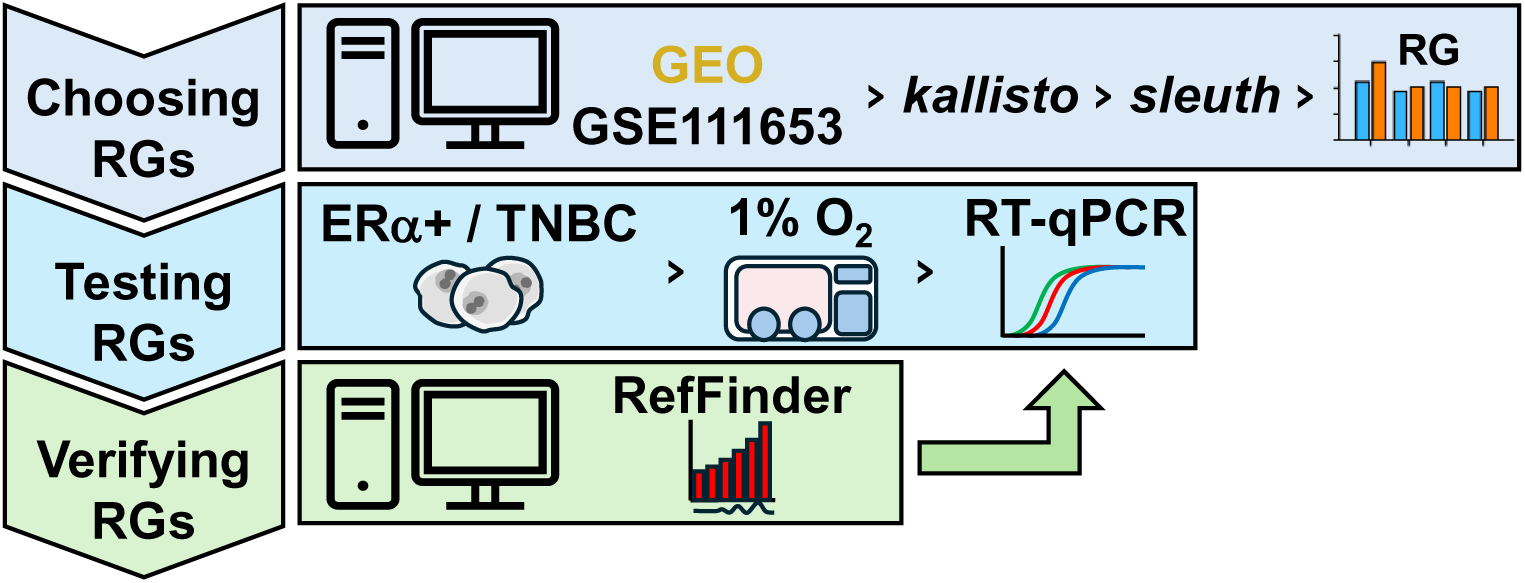

## Introduction

Breast cancer is the most common malignancy diagnosed worldwide. Due to advancements in therapeutic strategies and early detection, overall survival has greatly improved over the last few decades ^1^. However, approximately one third of diagnoses will result in death as a consequence of chemotherapeutic resistance, metastasis, or delayed presentation of treatment toxicities ^2–4^. Therefore, identifying novel molecular targets for therapeutic intervention is imperative. Current hormonal therapies targeting oestrogen receptor (ERα) activity have been effectively used to treat ERα positive (ERα+) Luminal A and Luminal B breast cancers, which account for ∼70% of diagnoses ^5^. However, breast cancers that co-express ERα and human epidermal growth factor receptor 2 (HER2), HER2 alone, or triple negative breast cancers (TNBC) which don’t express any hormone or growth factor receptors are more aggressive and tougher to treat. Furthermore, acquired or *de novo* resistance to ERα-targeting drugs is an additional barrier that further diminishes survival for women with ERɑ+ disease ^6^.

Solid tumours, including those of the breast, have regions of limited O_2_ availability (hypoxia) due to increased O_2_ consumption in rapidly dividing cancer cells, and inadequate perfusion and diffusion of O_2_ as cells outgrow local blood vessel supply ^7,8^. Hypoxia-inducible factors (HIF)-1α and HIF-2α accumulate in hypoxic cells and are key transcriptional regulators of the hypoxic response. HIF-α subunits are constitutively expressed, even when O_2_ is abundant. However, under physiological levels of O_2_, HIF-α proteins are rapidly degraded by the proteasome via a tightly regulated process involving prolyl hydroxylase domain (PHD) enzymes and von Hippel-Lindau protein (pVHL) ^9^. PHD enzymes use O_2_ as a catalytic substrate to hydroxylate HIF-α proteins, and are inhibited under hypoxic conditions; this in turn inhibits proteasomal degradation, and thus promotes accumulation of HIF-α subunits ^10^. Stabilised HIF-α subunits translocate into the nucleus whereby they form heterodimers with HIF-1β and bind to hypoxia response elements (HREs) present within promotors of target genes, thereby initiating transcription. In solid tumours, hypoxia and HIFs are recognised as important contributors to cancer progression and metastasis ^11^. Hypoxia has been shown to remodel the chromatin landscape of breast cancer cells to promote epithelial-to-mesenchymal transition (EMT) in a HIF-1α-dependent manor ^12^. Additionally, hypoxia has been linked to chemotherapy and radiotherapy resistance, and poor disease outcome ^13–15^.

To assess complex physiological changes occurring during hypoxia-mediated breast cancer progression and therapy resistance, reverse transcription - quantitative real-time polymerase chain reaction (RT-qPCR) is gold standard for accurately quantifying gene transcription and capturing dynamic changes in gene expression that may be serving as molecular drivers of advanced disease ^16^. A fundamental component of RT-qPCR is inclusion of reference genes (RGs) which act as internal controls for endogenous normalisation of measured target gene expression. RGs are selected on the basis of constitutive expression, and relative abundance not being altered by experimental conditions^17^. The substantial adjustment to the epigenome and transcriptome of cells that occur under hypoxic conditions render traditional RGs such as glycolytic enzymes *GAPDH* or *PGK1* redundant; despite this, comprehensive, systemic determination of RGs for hypoxia studies has yet to be performed ^18–21^.

We sought to fill this important knowledge gap by identifying RGs suitable for interrogating effects of hypoxia in breast cancer progression, using four widely used breast cancer cell lines representing both ERα+ Luminal A (MCF-7 and T-47D) and TNBC (MDA-MB-231 and MDA-MB-468) subtypes. We identified 10 RG candidates following analysis of a publicly available RNA-seq dataset ^22,23^. We then established a comprehensive investigation of candidates to determine RGs with the least variability in expression after being cultured in normoxia (20% O_2_), acute hypoxia (1% O_2_, 8 hours) or chronic hypoxia (1% O_2_, 48 hours).

RG candidates not abundantly expressed or associated with poor primer efficiencies were filtered out of the selection process. RGs were chosen by employing web-based RG tool RefFinder ^24,25^. Our findings identify *RPLP1,* or *RPLP1* in combination with *RPL27* as optimal RGs for analysis of hypoxia-mediated gene transcription in MCF-7, T-47D, MDA-MB-231 and MDA-MB-468 breast cancer cell lines. Our identification of *RPLP1* and *RPL27* as robust RGs in the context of hypoxic breast cancer cell lines provides a valuable resource for future studies investigating important transcriptional changes occurring during breast cancer progression.

## Materials and Methods

### Cell Culture

MCF-7, T-47D, MDA-MB-231 and MDA-MB-468 cell lines were used in this investigation. All breast cancer cell lines were maintained in DMEM (Gibco; S41966-029) supplemented with 5% foetal bovine serum (FBS; Gibco; 10270106) in a humidified Binder CO_2_ incubator at 37°C and 5% CO_2_. Cells were regularly tested for *Mycoplasma* by immunofluorescent visualisation of *Mycoplasma* DNA with DAPI ^26^. T-47D cells were provided by Dr. Andrew Holding (University of York) and MDA-MB-231 cells were a gift from Prof. Mustafa Djamgoz (Imperial College London). Both T-47D and MDA-MB-231 cell lines were authenticated by STR profiling. The MCF-7 and MDA-MB-468 cell lines were purchased from ATCC. For hypoxia culture, breast cancer cell lines were incubated in a humidified Baker Ruskinn InvivO_2_ oxygen workstation (37°C, 1% O_2_, 5% CO_2_) for 8 or 48 hours.

### Selection of RG candidates

High throughput RNA-seq datasets of 32 breast cancer cell lines cultured in 20% or 1% O_2_ for 24 hours are available from the NCBI Gene Expression Omnibus (GEO; Series Accession: GSE111653) ^22,23^. Using the University of York’s Viking 2 cluster, we recovered paired-end fastq files for hypoxic and normoxic MCF-7, T-47D, MDA-MB-231 and MDA-MB-468 breast cancer cells with *fastq-dump* (Supplementary Table S1). Low-quality reads were trimmed with *trimmomatic* (ILLUMINACLIP: TruSeq3-PE.fa:2:30:20 LEADING:3 TRAILING:3 SLIDINGWINDOW:4:15 MINLEN:36) and fastQC reports were generated with *fastQC*. Reads were pseudoaligned to the GRCh38.p14 annotation (release 111) and quantified using *kallisto.* Hierarchical Data Format (h5) files containing quantified reads for each experiment were input into RStudio (version 4.3.3). Here, quantified reads were aggregated on the gene level using *sleuth_prep (gene_mode* = TRUE) for differential analysis.

To determine relative stability across a selection of common RGs, and generate a shortlist of RG candidates, normalised reads in transcript per million (TPM) at common RGs in hypoxia and normoxia were assessed independently for each cell line. A shortlist of RG candidates was selected based on (i) the appearance of the RG in literature searches and/or (ii) had a calculated similarity (s) score of ≤ 0.3 between the 20% and 1% O_2_ conditions in at least two of the breast cancer cell lines. s was calculated by s = 1 - MIN(A,B) / MAX(A,B) (Microsoft Excel), where A is the read count value for a gene in 1% O_2_, B is the read count value for a gene in 20% O_2_, MIN refers to the smallest value between A and B and MAX determines the maximum value between A and B.

### RNA Isolation and cDNA Synthesis

Breast cancer cell lines were seeded to a density of 0.2 x10^6^ per well of a 6-well plate and were left for a minimum of 24 hours to adhere to the surface of wells before starting experiments. Each experiment was carried out with three biological replicates, consisting of three technical replicates. The experiment was laid out such that all samples from each normoxic or hypoxic timepoint were collected on the same day. At the experiment endpoint, cold QIAzol lysis reagent (QIAgen; 79306) was used to harvest RNA, as per manufacturer’s guidelines. Samples were rapidly collected in QIAzol, placed on ice and stored at −80°C before RNA extraction. For phase separation, phenol/chloroform extraction with isopropanol precipitation was carried out as previously described ^27^. To enhance nucleic acid extraction, GlycoBlue Coprecipitant (Invitrogen; AM9515) was included in the isolation protocol. Nucleic acid was re-suspended in 0.2 μm-filtered RNase-free water (Ambion; AM9937) and treated with DNase I (New England BioLabs; M0303S) to remove contaminating genomic DNA. RNA concentration and purity were measured using a NanoDrop™ One/OneC Microvolume UV-Vis Spectrophotometer (Thermo Fisher Scientific). RNA with an A260/280 of ≥ 2.0 was used. To ensure integrity, RNA was assessed by 1.5 % agarose gel electrophoresis in denaturing conditions.

RNA was reverse transcribed using SuperScript IV cDNA Synthesis Kit as per manufacturer’s instructions (Invitrogen; 18091050). The amount of total RNA was 1 μg. The reaction volume was 20 μl and consisted of 1 μl 0.1M DTT, 4 μl SSIV buffer, 1 μl RNAseOUT, 1 μl SSIV Enzyme, 1 μg of RNA in 11 μl of dH2O, 1 μl of random hexamer and 1 μl of 10 mM dNTP. Reactions were carried out on a Bioer LifePro thermocycler, comprising an initial step at 65°C for 05:00 (mm:ss), followed by 23°C for 10:00 (mm:ss), 55°C for 10:00 (mm:ss), 80°C for 10:00 (mm:ss) and then 4°C for 10:00 (mm:ss). cDNA samples were diluted to 5 ng / μl in 0.2 μm-filtered RNase-free water (Ambion; AM9937). A standard curve was prepared from pooled RNA from each biological replicate, and diluted to 20 ng / μl, 4 ng / μl, 0.8 ng / μl, 0.16 ng / μl and 0.032 ng / μl. Samples were stored at -30°C until further downstream analysis.

### RT-qPCR

RT-qPCR was performed using the QuantStudio™ 7 qPCR system (Thermo Fisher) in MicroAmp optical 384-well reaction plates (Applied Biosystems; 4309849) sealed with Expell™ optical sealing membranes (CAPP; 510400C). Technical reactions were performed in duplicate using 2X SYBR Green SuperMix (Applied Biosystems; 4385612). Each reaction mixture had a final working volume of 12 μl, containing 6 μl SuperMix, 1 μl 10 μM primer stock (Table 1) and 4 μl of 5 ng / μl cDNA. Primer sequences for *ACTB* ^28^, *RPL27* ^29^, *CCSER2* ^30^, *GUSB* ^31^, *TFRC* ^32,33^ and *CA9* ^34^ have been described before. For *OAZ1*, *TBP*, *RPL30*, *RPLP1*, *PGK1* and *EPAS1*, NCBI Primer BLAST was used to generate primer pair sequences that span the exon-exon junction with an amplicon size of between 70 and 200 bp and an optimal melting temperature of 60°C ± 3°C. All primer sequences were run through NCBI Primer BLAST to ensure no unintended gene targets could be amplified, but predicted transcript variants of the same gene were allowed. Primers were purchased from Integrated DNA Technologies.

**Table 1.**
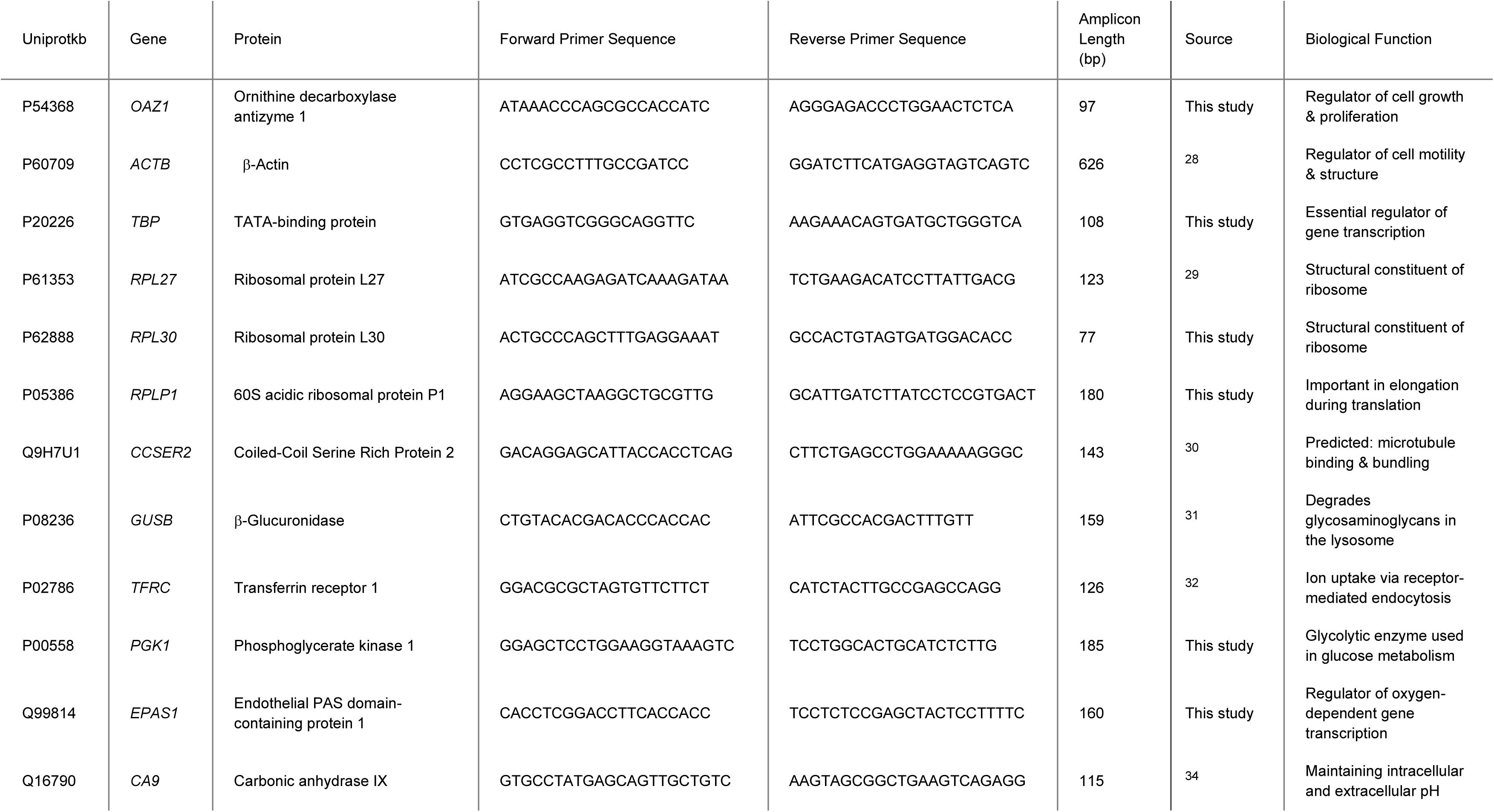
Primer sequences and protein features of reference genes and hypoxia responders.

For the standard curve, 80 ng, 16 ng, 3.2 ng, 0.64 ng and 0.128 ng of pooled cDNA was used. No-template reactions were included as a negative control. RT-qPCR cycling parameters comprised an initial denaturation step at 95°C for 01:35 (mm:ss), followed by 40 cycles of 00:03 (mm:ss) at 95°C and 00:30 (mm:ss) at 60°C. Melt curve analysis was carried out in the final cycle of the RT-qPCR by increasing the temperature from 60°C to 95°C at 0.1°C per second.

### Determining RG stability in breast cancer cell lines in normoxia, or acute or chronic hypoxia

Following RT-qPCR, reaction summaries were exported from ThermoFisher Design and Analysis Data Gallery and analysed in Microsoft Excel. A standard curve was used to calculate primer efficiency (PE) using the equation PE% = (10^(−1/*m*)^ − 1) ∗ 100 where *m* denotes the slope of the standard curve. Then, PE = *SUM*(*PE*%/100) + 1. Where PE was > 2.20 or, < 1.80, RG candidates were excluded from further analysis. Efficiency-corrected Ct values (CtE) were calculated using the equation CtE = *SUM*(*Ct* ∗ (*log*(*PE*)/*log*(2))). mRNA expression (mE) of normoxic reference genes was determined by mE = 10^((*CtE*−*a*)/*m*)^ where *a* refers to the Y intercept.

CtE values were supplied to the online tool RefFinder, for determination of most stable reference genes to be used in normoxic vs. hypoxic breast cancer cell lines (available at https://www.ciidirsinaloa.com.mx/RefFinder-master) ^24^. RefFinder is a comprehensive program which employs computational RG analysis tools geNorm ^35^, Normfinder ^36^, BestKeeper ^37^ and the comparative ΔCt method ^38^ to rank candidate RGs based on the ranking from each RG analysis tools.

### Validation of RGs

The change in gene expression of the HIF-regulated, hypoxia-induced target gene *CA9* was assessed using the 2^-ΔΔCt^ method ^39^, with the geometric mean of the recommended RG combination for comparative analysis between MCF-7, T-47D, MDA-MB-231 and MDA-MB-468 breast cancer cell lines used for normalisation. One-way ANOVA and Dunnett’s multiple comparisons were performed to assess significant fold-change in *CA9* expression following normalisation with the geometric mean of the recommended pair of RGs. Statistical analysis was performed using GraphPad Prism. Significance was reported where *p < 0.05*.

## Results

### Analysis of public RNA-seq dataset identifies 10 RG candidates

The aim of our study was to identify optimal RGs for investigations of normoxic vs. hypoxic ERα+ Luminal A (MCF-7 and T-47D) and TNBC (MDA-MB-231 and MDA-MB-469) cell lines. We selected cell lines based on widespread use in breast cancer research: MCF-7, T-47D and MDA-MB-231 represent more than two-thirds of cell lines used within such studies ^40^.

To address the need for robust RGs, we first utilised a publicly available RNA-seq dataset that investigated genome-wide transcriptional changes taking place in 32 breast cancer cell lines as a consequence of O_2_ deprivation ^22,23^. From the 30,187 genes quantified in selected ERα+ and TNBC cell lines, we were able to evaluate overall distance between individual datasets (Supplementary Figure S1) and responsiveness of hypoxia-regulated genes to ensure cell lines behaved as expected when cultured in the absence of O_2_.

Analysis demonstrated increased expression of *CA9, PGK1* and *VEGFA* in all cell lines, in response to hypoxic culture and in line with previous findings (Supplementary Figure S2) _20,41–43_. We also looked at *ARNT* (HIF-1β), *ARNT2* (HIF-2β), *EPAS1, HIF1A* and *HIF3A* expression in each cell line in normoxia and hypoxia (Supplementary Figure S3). Interestingly, we found *EPAS1*, the gene encoding HIF-2α, appeared to be relatively stable in expression in TNBC but not ERα+ cell lines (Supplementary Figure S3; Table 2).

**Table 2.** Similarity (s) score between hypoxic and normoxic RNA-sequencing reads of RG candidates.

| RG candidate | MCF-7 | T-47D | MDA-MB-231 | MDA-MB-468 |
| --- | --- | --- | --- | --- |
| *ACTB* | 0.23 | 0.08 | 0.11 | 0.30 |
| *CCSER2* | 0.39 | 0.04 | 0.15 | 0.00 |
| *EPAS1* | 0.33 | 0.40 | 0.07 | 0.03 |
| *GUSB* | 0.11 | 0.20 | 0.08 | 0.21 |
| *OAZ1* | 0.34 | 0.11 | 0.05 | 0.03 |
| *RPL27* | 0.24 | 0.12 | 0.06 | 0.10 |
| *RPL30* | 0.29 | 0.07 | 0.04 | 0.01 |
| *RPLP1* | 0.15 | 0.22 | 0.01 | 0.03 |
| *TBP* | 0.14 | 0.19 | 0.09 | 0.34 |
| *TFRC* | 0.39 | 0.27 | 0.32 | 0.03 |

Next, we interrogated read count stability of common RGs when ERα+ and TNBC cells were cultured in hypoxia or normoxia, to identify RG candidates that may be stable in expression in each cell line, regardless of O_2_ availability (Supplementary Figure S4, Supplementary Figure S5). From this, we generated a shortlist of 10 RG candidates (Table 2; Supplementary Figure S4). We initially selected candidates based on common use as RGs in breast cancer cell lines (e.g. *CCSER2* in MCF-7, T-47D, MDA-MB-231 and MDA-MB-468 cell lines), or as stable RGs in other models of hypoxia (e.g. *RPLP1* in hypoxic pre-conditioned human neural stem cells) ^44–46^, and further stratified candidates based on a calculated similarity score (*s*) which was used to determine how similar read counts are in genes from breast cancer cell lines cultured in 20% or 1% O_2_. Where *s* = 0, read counts are the same between the two conditions. A minimum threshold was established where *s* ≤ 0.30 in at least two of the cell lines, to be carried forward as an RG candidate.

Of the 10 RG candidates, ERα+ MCF-7 cells had the greatest variability in RG expression compared to T-47D and the TNBC cell lines, with *CCSER2, EPAS1, OAZ1* and *TFRC* exceeding the maximum threshold for RG candidate selection, achieving *s* scores of 0.39, 0.33, 0.34 and 0.39, respectively (Table 2). Additionally, when *s* was calculated across the transcriptome of each breast cancer cell line, MCF-7 also had the highest percentage of genes exceeding the maximum threshold set as a marker of stable gene expression (Supplementary Figure S6). *EPAS1* also responded positively to hypoxic culture in T-47D cells with an *s* score of 0.40, whereas no induction was observed in the TNBC cells.

However, *EPAS1* was the only RG candidate that exceeded the maximum threshold in T-47Ds. Furthermore, for MDA-MB-231 and MDA-MB-468 cell lines, only *TFRC* or *TBP* had altered expression following O_2_ deprivation, with *s* scores of 0.32 and 0.34, respectively. Remaining RG candidates *ACTB, GUSB, RPL27, RPL30* and *RPLP1* were stable in expression between the two conditions, in all cell lines (Table 2; Supplementary Figure S4).

### Assessment of RG candidate mRNA expression in normoxia confirms eight highly expressed genes

To demonstrate suitability of RG candidates, we sought to confirm RG expression in TNBC and ERα+ breast cancer cell lines cultured in normal O_2_ conditions. *ACTB* was expressed most highly among the breast cancer cell lines, but also showed greatest variation between biological replicates ranging from 8 ng / μl to 202 ng / μl in MCF-7 cells, and 30 ng / μl to 179 ng / μl in T-47D cells (Figure 1A). *EPAS1* was only amplified in one biological replicate in MDA-MB-231 and MDA-MB-468 cells, with mRNA levels of 103 ng / μl and 6 ng / μl, respectively (Figure 1C). Additionally, *TBP* did not have detectable levels of transcript in any cell lines (Figure 1J). These results are supported by the RNA-seq analysis (Supplementary Figure S3, Supplementary Figure S4), where TPM for *EPAS1* and *TBP* were among the lowest in expression in the breast cancer cell lines compared to other RG candidates. We therefore removed *TBP* and *EPAS1* from further investigation. The next lowest expressed RG was *CCSER2* which was expressed at 0.21, 0.31, 1.15 and 2.36 ng / μl in MCF-7, T-47D, MDA-MB-468 and MDA-MB-231 cell lysates, respectively (Figure 1B). The remaining RG candidates (*GUSB*, *OAZ1*, *RPL27*, *RPL30*, *RPLP1* and *TFRC*) and *PGK1* were more highly expressed in all cell lines (Figure 1D – Figure 1I, Figure 1K).

**Figure 1.**
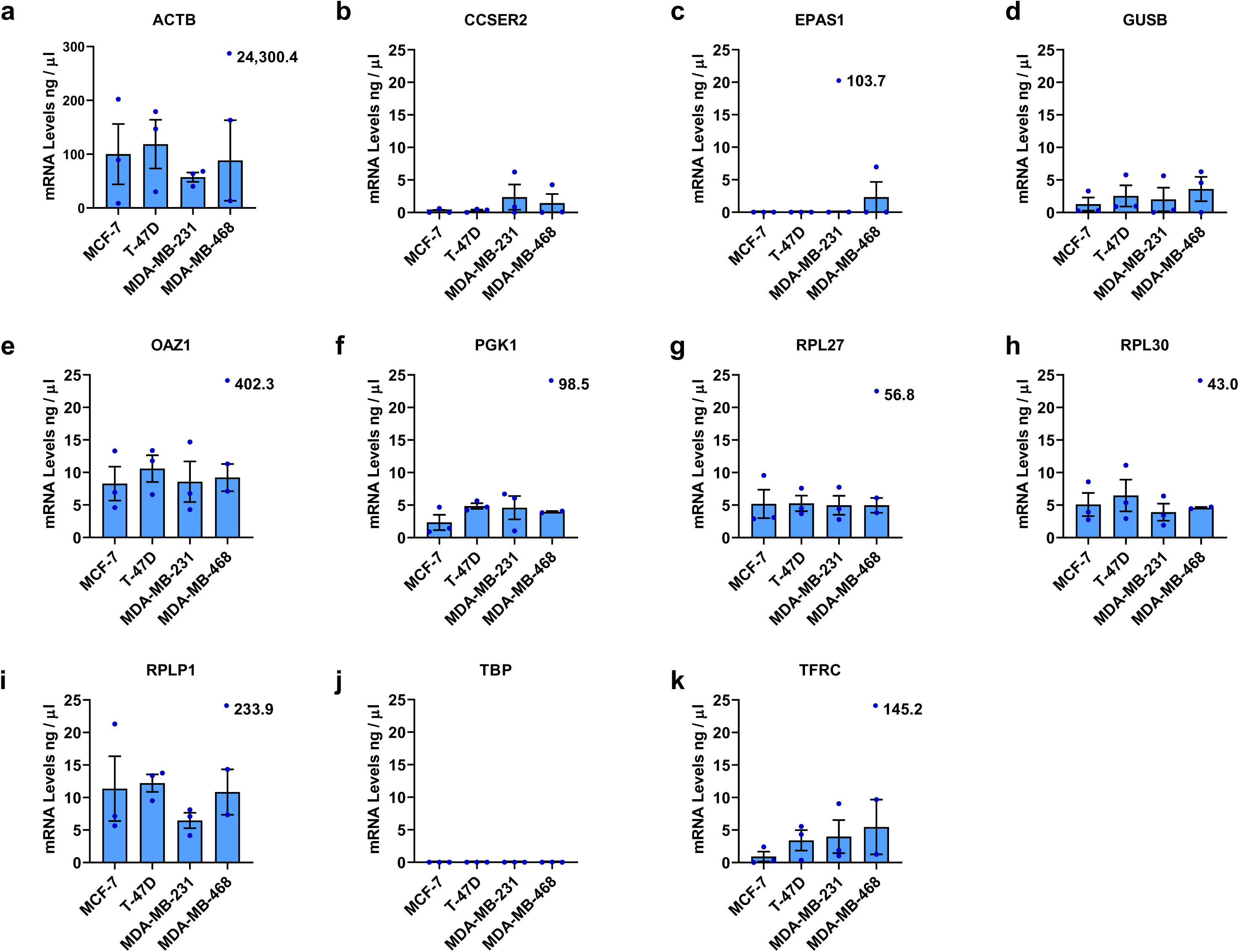
Expression of RG candidates in MCF-7, T-47D, MDA-MB-231 and MDA-MB-468 breast cancer cell lines cultured in 20% O_2_. Selected RG candidates (A) ACTB, (B) CCSER2, (C) EPAS1, (D) GUSB, (E) OAZ1, (F) PGK1, (G) RPL27, (H) RPL30, (I) RPLP1 (J) TBP and (K) TFRC were evaluated for mRNA expression in breast cancer cell lines cultured in normal conditions for 72 hours post seeding. Error bars are ± SEM. n = 3. Where there is an outlier, data point is displayed above the relevant box plot with mRNA expression value included.

### Evaluating RG expression in normoxic vs. hypoxic breast cancer cell lines filters out poor RG candidates and identifies robust RGs with the least variability in expression

Next, expression stability of RG candidates was investigated following breast cancer cell line culture in normoxia, or in hypoxia for 8 or 48 hours (Figure 2A; Supplementary Table S2). We also tested PEs from standard curves included in RT-qPCR experiments (Supplementary Table S3; Supplementary Figure S7 – Supplementary Figure S10). *ACTB, CCSER2* and *GUSB* displayed poor PE (Supplementary Table S3; *ACTB* mean 1.70, range 1.48 - 1.96; *CCSER2* mean 2.43, range 2.05 - 3.26; *GUSB* mean 2.22, range 2.03 - 2.50).

**Figure 2.**
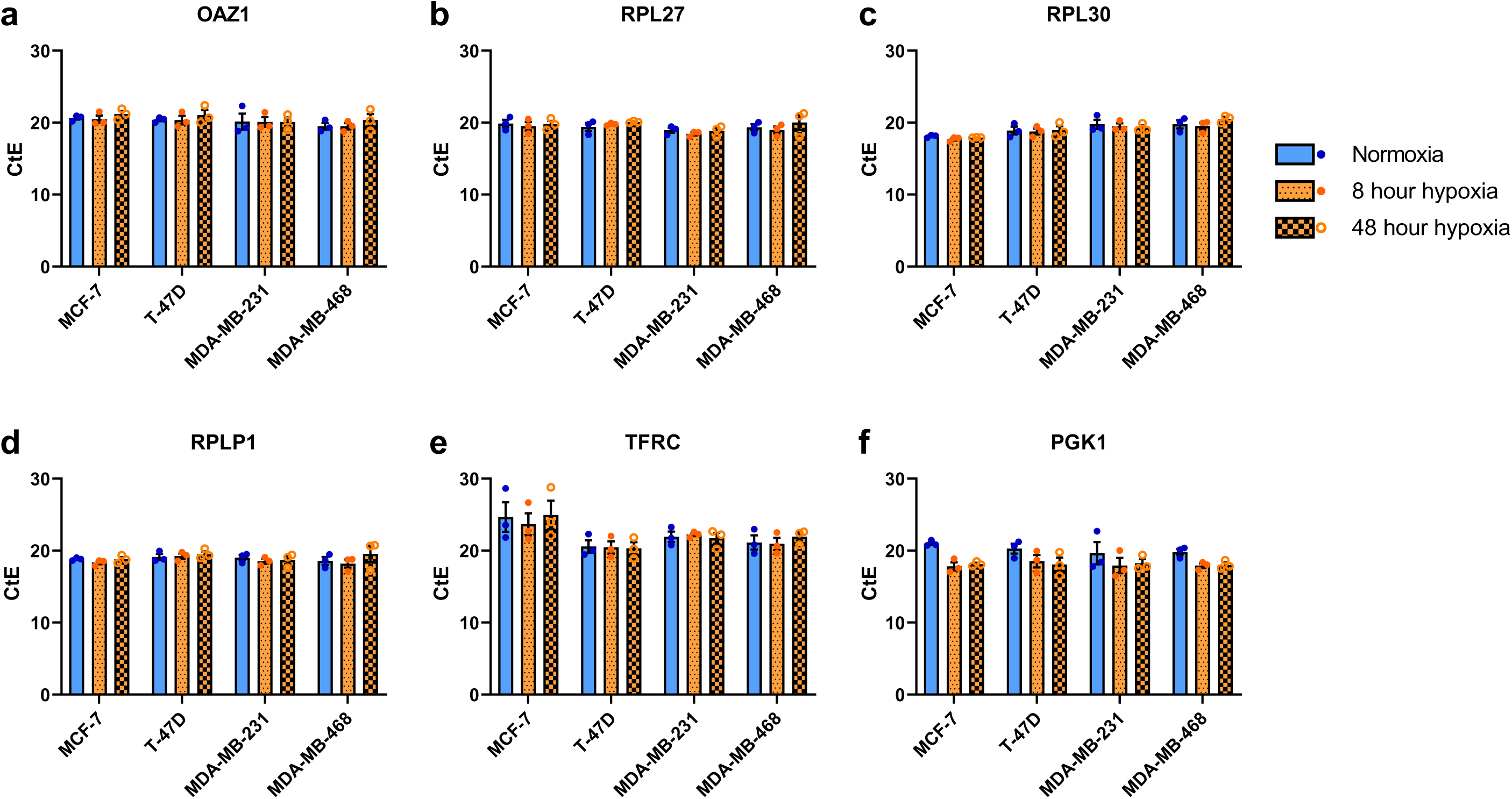
RG stability in breast cancer cell lines cultured in normoxia for 72 hours post-seeding, or normoxia and then hypoxia for 8 or 48 hours (total experimental time 72 hours post-seeding). RT-qPCR was used to determine the variance in gene expression of selected RG candidates: (A) *OAZ1*, (B) *RPL27*, (C) *RPL30*, (D) *RPLP1*, (E) *TFRC* and (F) *PGK1* following culture of MDA-MB-231, MDA-MB-468, MCF-7 or T-47D breast cancer cell lines in normoxia (blue bars, closed blue points - left) or hypoxia for 8 hours (orange bars, closed orange points - middle) or 48 hours (orange checkered bars, open orange points - right). Error bars are ± SEM

These RG candidates were therefore removed from downstream analysis. *OAZ1, RPL27, RPL30* and *RPLP1* were expressed at comparatively similar levels across all cell lines, and in each condition (Figure 2A - Figure 2D). *TFRC* shows inter-cell line stability when cultured in normoxia, or acute or chronic hypoxia. However, intra-cell line CtE was more varied. In particular, *TFRC* had higher CtE values in MCF-7 cells, which suggests this gene is not as highly expressed in MCF-7s compared to other breast cancer cell lines (Figure 2E). As predicted based on literature, *PGK1* CtE values decreased in all cell lines following hypoxic culture for 8 or 48 hours, which confers increased expression of *PGK1* in response to limited O_2_ supply (Figure 2F). This result is in line with previous observations of hypoxic induction of PGK1^20,45,47,48^.

We then submitted CtE values (Supplementary Table S2) of the five remaining RG candidates, *OAZ1, RPL27, RPL30, RPLP1* and *TFRC,* as well as hypoxia-responder *PGK1,* to RefFinder, with intent to rank RG candidates in order of expression stability across all cell lines in normoxia or acute or chronic hypoxia. RefFinder first employs GeNorm, NormFinder, BestKeeper and the comparative ΔCt method to independently rank RGs. Next, RefFinder assigns a weight to an individual gene based on RG performance in the prerequisite programs, and calculates the geometric mean of candidate weights to provide a final ranking of the most stable RGs ^24,25^. In all iterations of RG stability analysis across all cell lines, *PGK1* and *TFRC* were ranked 5th and 6th, respectively (Table 3). According to BestKeepeer and the comparative ΔCt method, *RPLP1* has the least variable inter- and intra-cell line expression in normoxic and hypoxic environments. *RPLP1* was also the highest ranked RG candidate by RefFinder (Figure 3A). Conversely, NormFinder ranked *OAZ1* as the best RG candidate, and placed *RPL27* and *RPLP1* as the second and third best RG candidates (Table 3). A benefit of GeNorm over the other programs is the additional assessment of the optimal number of RGs to use for accurate normalisation ^35^. For the study of hypoxia-mediated alterations in gene expression between MCF-7, T-47D, MDA-MB-231 and MDA-MB-468 breast cancer cell lines, GeNorm recommends the combined use of *RPL27* and *RPLP1*.

**Figure 3.**
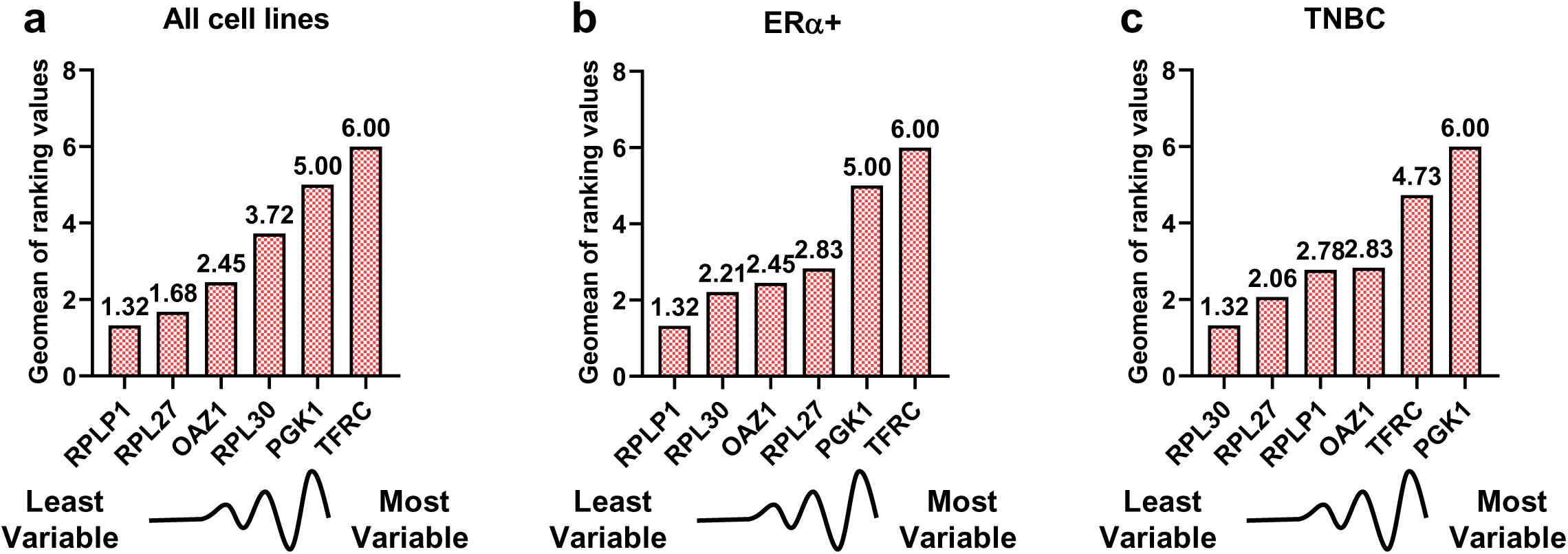
Geometric mean (Geomean) of ranking values for each RG candidate according to RefFinder. The final overall ranking of RG candidates was determined by RefFinder based on the geometric mean of the weights of each gene from GeNorm, NormFinder, BestKeeper and the comparative ΔCt method for (A) all breast cancer cell lines, (B) ERα+ breast cancer cell lines MCF-7 and T-47D and (C) TNBC cell lines MDA-MB-231 and MDA-MB-468.

**Table 3.** Summary of individual rankings of RG stability according to RefFinder, comparative ΔCt method, BestKeeper, NormFinder and GeNorm. Ranking was determined by providing all *CtE* values for each RG assessed in**, MCF-7, T-47D, MDA-MB-231** and **MDA-MB-468** breast cancer cell lines, in all conditions to RefFinder.

| Method | Ranking (1 = best - 6 = worst) |  |  |  |  |  |
| --- | --- | --- | --- | --- | --- | --- |
|  | 1 | 2 | 3 | 4 | 5 | 6 |
| $\Delta C_t$ | RPLP1 | RPL27 | OAZ1 | RPL30 | PGK1 | TFRC |
| BestKeeper | RPLP1 | RPL27 | OAZ1 | RPL30 | PGK1 | TFRC |
| Normfinder | OAZ1 | RPL27 | RPLP1 | RPL30 | PGK1 | TFRC |
| GeNorm | RPL27 \| RPLP1 |  | RPL30 | OAZ1 | PGK1 | TFRC |
| RefFinder recommended comprehensive ranking | RPLP1 | RPL27 | OAZ1 | RPL30 | PGK1 | TFRC |

We next identified optimal RGs to be used for RT-qPCR of hypoxic breast cancer cell lines following stratification into breast cancer subtypes. When CtE values from ERα+ breast cancer cell lines were supplied, *RPLP1* was again ranked top RG candidate with the least variability in expression, according to RefFinder, BestKeepeer and the comparative ΔCt method (Table 4, Figure 3B). As in all cell lines, Normfinder suggests *OAZ1* to be the optimal RG to use when investigating hypoxic induction of genes of interest in the ERα+ Luminal A breast cancer group. GeNorm recommends the combined use of *RPLP1* and *RPL30*, instead of *RPL27* as previously put forward for all cell lines. *PGK1* and *TFRC* were ranked as the least stable RGs in all outputs as before. For the TNBC group, *RPL30* was placed first by all programs (Table 5, Figure. 3C), apart from GeNorm which recommended *RPL27* and *RPLP1,* the same as for all breast cancer cell lines. Computational analysis of individual cell lines cultured in normoxia, and acute or chronic hypoxia was also performed. Here, GeNorm identifies *RPLP1* and *RPL27* as the least variable and most suitable RGs for MDA-MB-231 or MDA-MB-468 cell lines, but *RPL30* is ranked as the least variable single RG by RefFinder in both TNBC models. *RPLP1* and *RPL30* are the least variable and most suitable RGs for the T-47D or MCF-7 cell lines (Supplementary Tables S4-S7, Supplementary Figure S11).

**Table 4.** Summary of individual rankings of RG stability according to RefFinder, comparative ΔCt method, BestKeeper, NormFinder and GeNorm. Ranking was determined by providing *CtE* values for each RG assessed in **MCF-7** and **T-47D ERα+** breast cancer cell lines, in all conditions to RefFinder.

| Method | Ranking (1 = best - 6 = worst) |  |  |  |  |  |
| --- | --- | --- | --- | --- | --- | --- |
|  | 1 | 2 | 3 | 4 | 5 | 6 |
| $\Delta C_t$ | RPLP1 | RPL30 | OAZ1 | RPL27 | PGK1 | TFRC |
| BestKeeper | RPLP1 | RPL27 | RPL30 | OAZ1 | PGK1 | TFRC |
| Normfinder | OAZ1 | RPL27 | RPLP1 | RPL30 | PGK1 | TFRC |
| GeNorm | RPL30 \| RPLP1 |  | OAZ1 | RPL27 | PGK1 | TFRC |
| RefFinder recommended comprehensive ranking | RPLP1 | RPL30 | OAZ1 | RPL27 | PGK1 | TFRC |

**Table 5.**
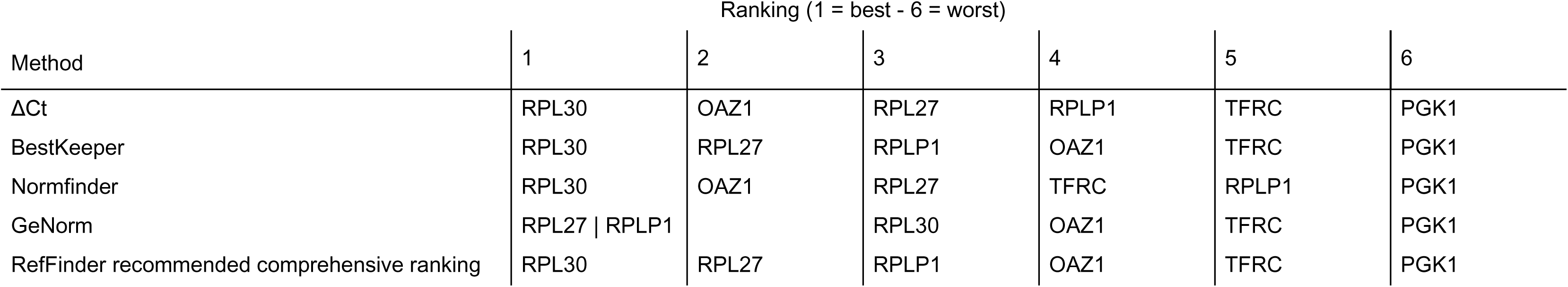
Summary of individual rankings of RG stability according to RefFinder, comparative ΔCt method, BestKeeper, NormFinder and GeNorm. Ranking was determined by providing all *CtE* values for each RG assessed in **MDA-MB-231** and **MDA-MB-468** TNBC cell lines, in all conditions to RefFinder.

| Method | Ranking (1 = best - 6 = worst) |  |  |  |  |  |
| --- | --- | --- | --- | --- | --- | --- |
|  | 1 | 2 | 3 | 4 | 5 | 6 |
| $\Delta C_t$ | RPL30 | OAZ1 | RPL27 | RPLP1 | TFRC | PGK1 |
| BestKeeper | RPL30 | RPL27 | RPLP1 | OAZ1 | TFRC | PGK1 |
| Normfinder | RPL30 | OAZ1 | RPL27 | TFRC | RPLP1 | PGK1 |
| GeNorm | RPL27 \| RPLP1 |  | RPL30 | OAZ1 | TFRC | PGK1 |
| RefFinder recommended comprehensive ranking | RPL30 | RPL27 | RPLP1 | OAZ1 | TFRC | PGK1 |

### RPLP1 and RPL27 are suitable RGs for normalising gene expression in normoxic vs. hypoxic ERα+ and TNBC cell lines

Following identification of optimal RGs, we aimed to evaluate combined use of *RPLP1* and *RPL27* for normalisation of gene transcription in normoxic and hypoxic breast cancer cell lines. We assessed upregulation of hypoxia-induced *CA9* in each breast cancer cell line cultured in normoxia or hypoxia for 8 or 48 hours. The geometric mean of *RPLP1* and *RPL27* was used to normalise *CA9* CtE values, before fold change induction (2^-ΔΔCt^) of *CA9* was calculated ^39^. Expression (CtE) of *RPLP1* and *RPL27* in MCF-7 (19.4 ± 0.4 SD), T-47D (19.7 ± 0.5 SD), MDA-MB-231 (18.8 ± 0.5 SD) and MDA-MB-468 (19.6 ± 0.9 SD) cells were consistent, regardless of environmental O_2_ (Figure 4A). Conversely, all cell lines demonstrate significant induction of *CA9* following hypoxic culture (Figure 4B). In MCF-7 cells, *CA9* was increased 470-fold after chronic exposure to a hypoxic environment. For T-47Ds, acute and chronic hypoxia induced a 42- and 109-fold increase in *CA9* expression, respectively. After 8 hours of hypoxic culture, MDA-MB-231s instigated a moderate but significant 9-fold induction, and for MDA-MB-468s a 17-fold increase in *CA9* expression was seen following 48 hours of hypoxic culture. Importantly, *RPLP1* and *RPL27* were similarly expressed in each cell line, in each condition. Thus, combination of *RPLP1* and *RPL27* as RGs is suitable for normalising gene expression in normoxic and hypoxic breast cancer cell lines.

**Figure 4.**
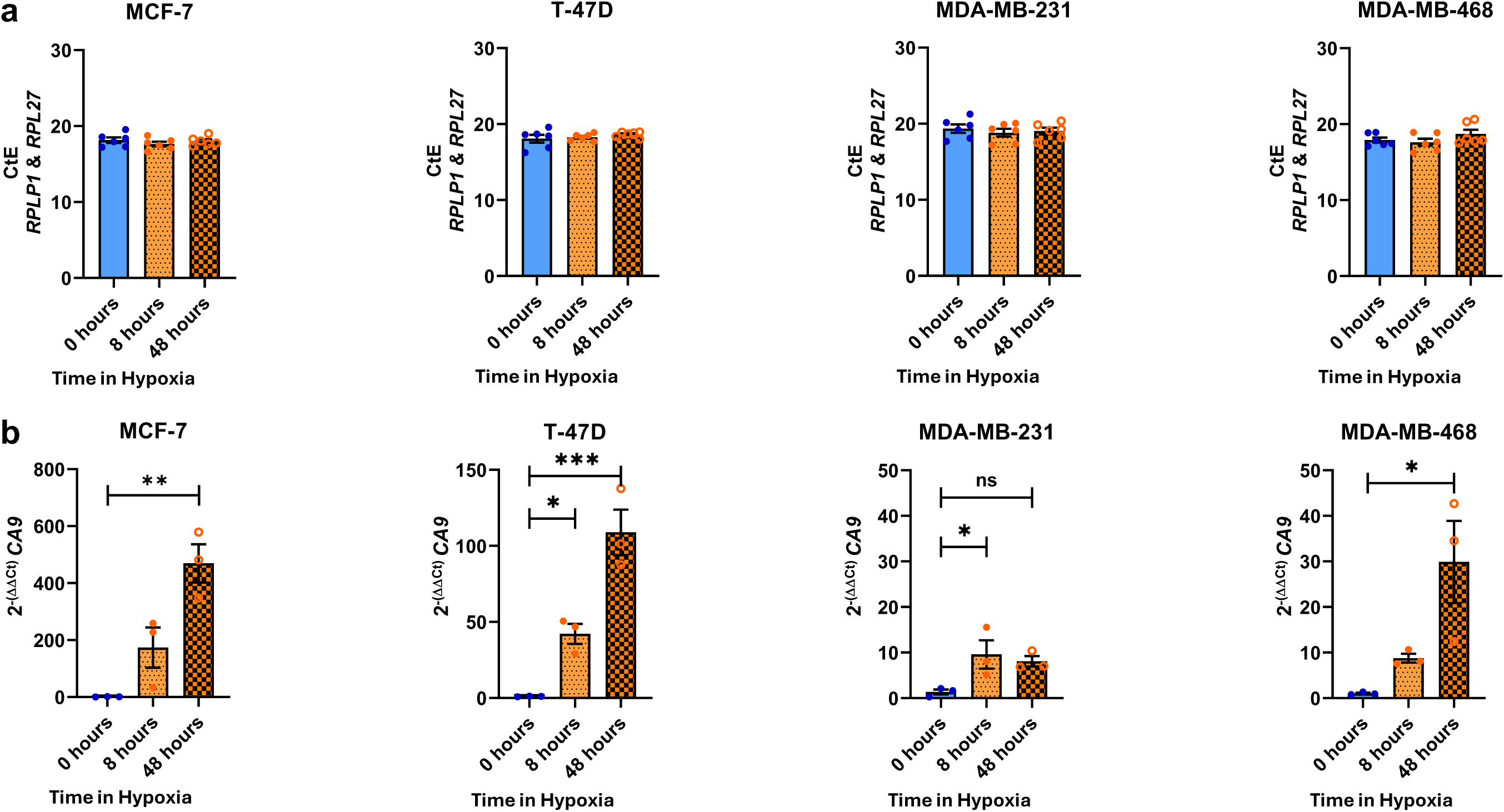
RG expression level stability and hypoxic induction of *CA9* in four breast cancer cell lines cultured in normoxia, or hypoxia for 8 or 48 hours. (A) *RPL27* (n = 3) and *RPLP1* (n = 3) expression was determined by RT-qPCR. Raw CtE values for triplicate biological replicates of the two RGs (n = 6) in MDA-MB-231, MDA-MB-468, T-47D and MCF-7 breast cancer cell lines are shown. Error bars are geometric mean ± geometric SD. (B) Expression of *CA9* was assessed in MCF-7, T-47D, MDA-MB-231 and MDA-MB-468 breast cancer cell lines following culture in normoxia (20% O_2_, “0 hours”) or hypoxia (1% O_2_) for 8 or 48 hours. Changes in *CA9* expression were determined by the 2^-ΔΔCt^ method, using the geometric mean of RGs *RPLP1* and *RPL27* for normalisation (A). One-way ANOVA with Dunnett’s multiple comparisons was employed to investigate significant fold change in gene expression relative to normoxic control. \**p= < 0.05,* \*\**p = < 0.01,* \*\*\**p = < 0.001*. Error bars are ± SEM. n = 3

## Discussion

The use of RT-qPCR for investigating gene transcription has been customary practice in labs since quantitative PCR was first discussed by Higuchi *et al.* in 1993 ^49^. While RT-qPCR is the gold standard for quantifying mRNA expression and understanding mechanisms involved in altered gene transcription, interpretation of gene expression is dependent on appropriate use of internal controls as a means of normalisation ^50^. Common RGs previously deemed stable in expression include *GAPDH, ACTB, PGK1* and *18S rRNA,* which have subsequently been shown to have variation in abundance across different experimental conditions, emphasising the notion that there is no such thing as an RG that works for all investigations ^51^. Indeed, in the context of cellular hypoxia, *ACTB* is under the influence of insufficient O_2_ supply, as are *GAPDH* and *PGK1* which are specifically regulated by the activity of HIF-1α _19–21,48_. Thus, when looking to identify novel therapeutic targets to combat hypoxia-induced therapy resistance for breast cancer patients, suitable RGs need to be selected prior to RT-qPCR investigation of genes of interest, so hypoxia-induced alterations in RG expression do not obscure novel and important biological findings.

To meet the demand for robust RGs for investigations of hypoxic ERα+ and TNBC cell lines, we carried out a comprehensive investigation combining bioinformatic analysis of publicly available RNA-seq datasets to select 10 RG candidates, RT-qPCR of candidates to assess expression levels and variability, and utilisation of the online RG tool RefFinder to ensure the most suitable RGs were selected. The 10 RG candidates we identified included genes that are generally considered RGs (*ACTB, RPL30, RPLP1, GUSB, TBP* and *TFRC*), and novel RGs (*OAZ1, RPL27, CCSER2*, and *EPAS1*) ^29,44–46,52,53^. When CtEs of our candidates were supplied to RG selection tools, it is perhaps unsurprising that constituents of the ribosome (*RPLP1, RPL27* and *RPL30*) which are abundantly and consistently expressed in human tissues were selected as the optimal RGs with the least variability in expression in breast cancer cell lines cultured in normoxia, or acute or chronic hypoxia ^54–56^. This is supported by the observation that breast cancer cells can bypass hypoxia-mediated inhibition of protein synthesis through gene silencing of 4E-BP1, eEF2 kinase or tuberous sclerosis complex 2 (TSC2), maintaining a continuous requirement of translational machinery ^57^.

Throughout our study, we have chosen to include the process of RG candidate deselection, based on assessment of gene expression and primer efficiencies, as it is important to understand peripheral results which impact the quality of data interpretation.

Thus, for full transparency of our RG selection process, we have shown negative filtration of poor candidates as well as positive selection of stable candidates. To ensure precision in normalising expression of genes of interest, we recommend including two RGs in RT-qPCR studies, as use of a single RG for normalising gene expression may result in erroneous interpretation, whereas inclusion of two RGs should ensure accurate normalisation of target gene abundance ^58,59^.

With respect to selection of our 10 RG candidates, the RNA-seq dataset used to curate the shortlist was limited by a single replicate for each cell line in each condition being available for analysis ^22,23^. The original study is an impressive investigation into the molecular portrait of hypoxia spanning 32 breast cancer cell lines and for the purpose of our study, provided a meaningful starting point for selecting and determining approximate stability of RG candidates. However, we acknowledge that for our interest in identifying stable RGs, the RNA-seq dataset alone would not be sufficient to draw robust conclusions about the optimal RGs to use in hypoxic breast cancer studies involving RT-qPCR.

A limitation of our study is identification of ribosomal proteins as suitable RGs may only be applicable to those wishing to capture hypoxia-induced changes in gene expression in MCF-7, T-47D, MDA-MB-231 and/or MDA-MB-468 breast cancer cell lines. How our results translate to other breast cancer cell lines, or indeed patient samples, remains unclear. Cell lines representing the same disease model often display variation in response to environmental or experimental conditions and have unique gene expression signatures and molecular portraits ^40^. This is exemplified in MCF-7 and T-47D cell lines, where 17β-oestradiol has been shown to confer disparate changes in gene expression between the two models of Luminal A breast cancer, despite both cell lines being driven by ERα activity ^60^. For patient derived samples, the answer to identifying suitable RGs for RT-qPCR is more unclear, due to the complexity of individuality between patients, heterogeneity of cell types within the tumour microenvironment and uneven distribution of hypoxia observed throughout tumours. Cancer grade at diagnosis, and samples coming from secondary metastatic sites will also require further optimisation of RGs. Indeed, patterns of dysregulated ribosomal protein expression has been observed in human tissues, primary cell lines and tumours ^61^. Thus, careful identification of suitable RGs for such studies needs to be implemented prior to carrying out the experiment, and perhaps consideration of including a greater number of RGs (3 – 5 for more complex tissue samples) would narrow variability and allow more accurate normalisation in such instances ^35^. Nonetheless, we have outlined a robust strategy for selection of suitable RGs that can be implemented to a broad range of studies wishing to identify important transcriptional aberrations acting as drivers of breast cancer progression.

In conclusion, we have carried out a comprehensive investigation to identify the most suitable RGs with the least variability in their expression, which can be used in RT-qPCR studies of breast cancer cell lines cultured in normoxia or hypoxia. We used robust computational RG selection programs following stringent criteria of RG candidates and recommend the inclusion of *RPLP1* and *RPL27* in RT-qPCR studies as internal controls for accurate interpretation of gene expression results. Furthermore, this important result provides the means to assess the impact of hypoxia within breast cancer development and progression.

## Data Availability

Code and supporting data are available from https://zenodo.org/doi/10.5281/zenodo.13166160

## Supplementary Data Statement

Supplementary Data are available at NAR online

## Acknowledgements

The Viking cluster was used during this project, which is a high-performance compute facility provided by the University of York. We are grateful for computational support from the University of York, IT Services, and the Research IT team. The authors would also like to extend their gratitude to the Genomics Lab in the University of York Bioscience Technology Facility for their expert support.

## Funding

This work was supported by a studentship from the Biotechnology and Biological Sciences Research Council BB/T007222/1 to JRM, BB/V000071/1 to ANH and BB/Y513970/1 to WJB, Royal Society RGS\R2\202120 to ANH and KSB, Kay Kendall Leukaemia Fund KKL1377 to KSB and Medical Research Council (MR/X018067/1) to WJB, KSB and ANH.

## Conflicts of Interest

The authors declare no conflicts of interest

## Notes

https://zenodo.org/doi/10.5281/zenodo.13166160

### Competing Interest Statement

The authors have declared no competing interest.

### Summary of Updates

New version corrects some minor typographic errors.

