## Supplementary Data for "Identification of robust RT-qPCR reference genes for studying changes in gene expression in response to hypoxia in breast cancer cell lines"

Supplementary Table S1. SRA accession numbers of hypoxic (1% $O_2$, 24 hours) and normoxic breast cancer cell lines from BioProject PRJNA437670, series GSE111653

| SRA Run Accession number | Breast Cancer Cell Line | Breast Cancer Subtype | Environmental Oxygen Status |
|---|---|---|---|
| SRR6822831 | MCF-7 | Luminal A | Hypoxic |
| SRR6822832 | | | Normoxic |
| SRR6822837 | MDA-MB-231-PSOC | Basal B | Hypoxic |
| SRR6822838 | | | Normoxic |
| SRR6822841 | MDA-MB-468 | Basal A | Hypoxic |
| SRR6822842 | | | Normoxic |
| SRR6822857 | T-47D | Luminal A | Hypoxic |
| SRR6822858 | | | Normoxic |

Supplementary Table S2. Efficient-corrected Ct values (*CtE*) of RGs following RT-qPCR of cell lysates from normoxic, acutely hypoxic or chronically hypoxic culturing.

| | *OAZ1* | *RPL27* | *RPL30* | *RPLP1* | *TFRC* | *PGK1* |
|---|---|---|---|---|---|---|
| MDA-MB-231 | 18.80 | 19.43 | 19.21 | 19.48 | 20.76 | 18.42 |
| | 19.28 | 18.28 | 19.04 | 18.26 | 21.73 | 17.78 |
| | 22.31 | 19.11 | 21.02 | 19.28 | 23.26 | 22.70 |
| | 19.50 | 18.64 | 19.11 | 19.06 | 21.82 | 17.29 |
| | 19.33 | 18.67 | 19.07 | 18.54 | 22.28 | 16.47 |
| | 21.42 | 18.00 | 20.29 | 17.95 | 22.59 | 20.00 |
| | 18.96 | 19.43 | 18.93 | 19.38 | 20.24 | 17.54 |
| | 20.19 | 19.09 | 18.96 | 19.07 | 22.66 | 17.79 |
| | 21.11 | 17.97 | 19.92 | 17.60 | 22.22 | 19.38 |
| MDA-MB-468 | 18.81 | 20.04 | 20.56 | 19.46 | 20.81 | 20.10 |
| | 19.28 | 19.47 | 18.66 | 18.63 | 19.59 | 18.93 |
| | 20.37 | 18.53 | 20.08 | 17.59 | 22.96 | 20.36 |
| | 18.75 | 19.13 | 20.03 | 18.78 | 20.65 | 18.04 |
| | 19.52 | 19.68 | 18.56 | 18.74 | 19.64 | 17.31 |
| | 20.19 | 18.08 | 19.93 | 17.08 | 22.52 | 18.35 |
| | 18.86 | 20.88 | 20.67 | 20.63 | 20.82 | 17.80 |
| | 21.84 | 21.25 | 20.92 | 20.73 | 22.60 | 18.66 |
| | 20.29 | 17.94 | 19.94 | 17.24 | 22.41 | 17.50 |
| MCF-7 | 20.94 | 18.95 | 17.89 | 18.82 | 21.82 | 21.45 |
| | 20.19 | 20.75 | 18.16 | 19.00 | 23.54 | 20.72 |
| | 20.78 | 19.92 | 18.27 | 18.60 | 28.63 | 20.81 |
| | 21.51 | 18.47 | 17.92 | 18.60 | 21.61 | 18.87 |
| | 19.89 | 20.48 | 17.86 | 18.52 | 22.68 | 17.53 |
| | 20.00 | 19.54 | 17.34 | 17.90 | 26.67 | 17.00 |
| | 21.92 | 18.99 | 17.96 | 19.35 | 22.09 | 18.49 |
| | 20.42 | 20.58 | 17.88 | 18.73 | 23.96 | 18.04 |
| | 21.20 | 19.74 | 17.91 | 18.26 | 28.79 | 17.83 |
| T-47D | 20.70 | 18.31 | 18.79 | 18.61 | 19.45 | 20.65 |
| | 20.00 | 19.77 | 17.98 | 18.79 | 22.29 | 18.94 |
| | 20.46 | 20.11 | 19.89 | 19.94 | 19.98 | 21.22 |
| | 21.48 | 19.94 | 19.46 | 19.77 | 20.17 | 19.95 |
| | 20.07 | 19.58 | 18.01 | 18.57 | 22.02 | 16.95 |
| | 19.46 | 19.77 | 18.81 | 19.35 | 19.11 | 18.68 |
| | 22.39 | 20.22 | 20.00 | 20.26 | 20.24 | 19.75 |
| | 20.67 | 20.07 | 18.09 | 18.96 | 21.80 | 16.50 |
| | 20.02 | 19.97 | 18.69 | 19.30 | 18.92 | 17.97 |
| Mean | 20.30 | 19.41 | 19.05 | 18.86 | 22.04 | 18.77 |
| SE | 0.17 | 0.14 | 0.17 | 0.14 | 0.37 | 0.25 |
| SD | 1.00 | 0.87 | 1.00 | 0.83 | 2.23 | 1.49 |
| CV | 0.05 | 0.04 | 0.05 | 0.04 | 0.10 | 0.08 |

Supplementary Table S3: Primer Efficiencies of RG primers

| | *ACTB* | *CCSER2* | *GUSB* | *OAZ1* | *PGK1* | *RPL27* | *RPL30* | *RPLP1* | *TFRC* |
|---|---|---|---|---|---|---|---|---|---|
| MDA-MB-231 | 1.88 | 2.05 | 2.07 | 2.01 | 1.99 | 2.01 | 2.01 | 2.01 | 2.04 |
| | 1.64 | 2.19 | 2.23 | 1.99 | 1.94 | 1.97 | 2.05 | 1.96 | 2.18 |
| | 1.74 | 2.49 | 2.29 | 2.05 | 2.07 | 2.04 | 2.09 | 1.99 | 2.12 |
| MDA-MB-468 | 1.96 | 2.08 | 2.03 | 2.02 | 1.98 | 2.02 | 2.06 | 1.98 | 2.01 |
| | 1.73 | 3.26 | 2.44 | 2.01 | 1.99 | 2.04 | 2.02 | 1.96 | 2.06 |
| | 1.51 | 2.39 | 2.50 | 1.99 | 1.96 | 1.99 | 2.04 | 1.90 | 2.21 |
| MCF-7 | 1.95 | 2.16 | 2.05 | 2.03 | 2.02 | 2.01 | 2.03 | 1.98 | 2.07 |
| | 1.59 | 2.50 | 2.27 | 2.02 | 2.02 | 2.08 | 2.06 | 1.99 | 2.29 |
| | 1.48 | 2.99 | 2.33 | 2.03 | 1.97 | 2.01 | 2.04 | 1.93 | 2.63 |
| T-47D | 1.87 | 2.23 | 2.04 | 2.04 | 1.99 | 2.05 | 2.04 | 1.98 | 2.06 |
| | 1.49 | 2.52 | 2.24 | 2.00 | 1.95 | 2.09 | 2.00 | 1.96 | 2.42 |
| | 1.55 | 2.30 | 2.19 | 1.97 | 1.98 | 2.04 | 2.04 | 1.99 | 2.09 |
| Mean | 1.70 | 2.43 | 2.22 | 2.01 | 1.99 | 2.03 | 2.04 | 1.97 | 2.18 |
| SE | 0.03 | 0.06 | 0.02 | 0.00 | 0.01 | 0.01 | 0.00 | 0.00 | 0.03 |
| SD | 0.17 | 0.35 | 0.15 | 0.02 | 0.03 | 0.03 | 0.02 | 0.03 | 0.18 |
| CV | 0.10 | 0.14 | 0.07 | 0.01 | 0.02 | 0.02 | 0.01 | 0.01 | 0.08 |

Supplementary Table S4. Summary table to show individual rankings of RG stability according to RefFinder, the comparative ΔCt method, BestKeeper, NormFinder or GeNorm. Ranking was determined by providing all CtE values for each RG assessed in **MCF-7** breast cancer cell lines, in all conditions to RefFinder.

| | Ranking (1 = best - 6 = worst) | | | | | |
|---|---|---|---|---|---|---|
| Method | 1 | 2 | 3 | 4 | 5 | 6 |
| ΔCt | *RPL30* | *RPLP1* | *OAZ1* | *RPL27* | *PGK1* | *TFRC* |
| BestKeeper | *RPL30* | *RPLP1* | *OAZ1* | *RPL27* | *PGK1* | *TFRC* |
| Normfinder | *RPL30* | *RPLP1* | *RPL27* | *OAZ1* | *PGK1* | *TFRC* |
| GeNorm | *RPL30* \| *RPLP1* | | *OAZ1* | *RPL27* | *PGK1* | *TFRC* |
| RefFinder recommended comprehensive ranking | *RPL30* | *RPLP1* | *OAZ1* | *RPL27* | *PGK1* | *TFRC* |

Supplementary Table S5. Summary table to show individual rankings of RG stability according to RefFinder, the comparative ΔCt method, BestKeeper, NormFinder or GeNorm. Ranking was determined by providing all CtE values for each RG assessed in **T-47D** breast cancer cell lines, in all conditions to RefFinder.

| | Ranking (1 = best - 6 = worst) | | | | | |
|---|---|---|---|---|---|---|
| Method | 1 | 2 | 3 | 4 | 5 | 6 |
| ΔCt | *RPLP1* | *RPL30* | *RPL27* | *OAZ1* | *PGK1* | *TFRC* |
| BestKeeper | *RPL27* | *RPLP1* | *RPL30* | *OAZ1* | *TFRC* | *PGK1* |
| Normfinder | *RPLP1* | *RPL30* | *OAZ1* | *RPL27* | *PGK1* | *TFRC* |
| GeNorm | *RPL30* \| *RPLP1* | | *RPL27* | *OAZ1* | *PGK1* | *TFRC* |
| RefFinder recommended comprehensive ranking | *RPLP1* | *RPL30* | *RPL27* | *OAZ1* | *PGK1* | *TFRC* |

Supplementary Table S6. Summary table to show individual rankings of RG stability according to RefFinder, the comparative ΔCt method, BestKeeper, NormFinder or GeNorm. Ranking was determined by providing all CtE values for each RG assessed in **MDA-MB-231** breast cancer cell lines, in all conditions to RefFinder.

| | Ranking (1 = best - 6 = worst) | | | | | |
|---|---|---|---|---|---|---|
| Method | 1 | 2 | 3 | 4 | 5 | 6 |
| *ΔCt* | *RPL30* | *OAZ1* | *TFRC* | *RPL27* | *RPLP1* | *PGK1* |
| *BestKeeper* | *RPL27* | *RPLP1* | *RPL30* | *TFRC* | *OAZ1* | *PGK1* |
| *Normfinder* | *RPL30* | *OAZ1* | *TFRC* | *RPL27* | *RPLP1* | *PGK1* |
| *GeNorm* | *RPL27* \| *RPLP1* | | *RPL30* | *TFRC* | *OAZ1* | *PGK1* |
| *RefFinder recommended comprehensive ranking* | *RPL30* | *RPL27* | *RPLP1* | *OAZ1* | *TFRC* | *PGK1* |

Supplementary Table S7. Summary table to show individual rankings of RG stability according to RefFinder, the comparative ΔCt method, BestKeeper, NormFinder or GeNorm. Ranking was determined by providing all CtE values for each RG assessed in **MDA-MB-468** breast cancer cell lines, in all conditions to RefFinder.

| | Ranking (1 = best - 6 = worst) | | | | | |
|---|---|---|---|---|---|---|
| Method | 1 | 2 | 3 | 4 | 5 | 6 |
| ΔCt | *RPL30* | *RPL27* | *OAZ1* | *RPLP1* | *PGK1* | *TFRC* |
| BestKeeper | *RPL30* | *OAZ1* | *PGK1* | *RPL27* | *RPLP1* | *TFRC* |
| Normfinder | *RPL30* | *OAZ1* | *RPL27* | *PGK1* | *RPLP1* | *TFRC* |
| GeNorm | *RPL27* \| *RPLP1* | | *RPL30* | *OAZ1* | *PGK1* | *TFRC* |
| RefFinder recommended comprehensive ranking | *RPL30* | *RPL27* | *OAZ1* | *RPLP1* | *PGK1* | *TFRC* |

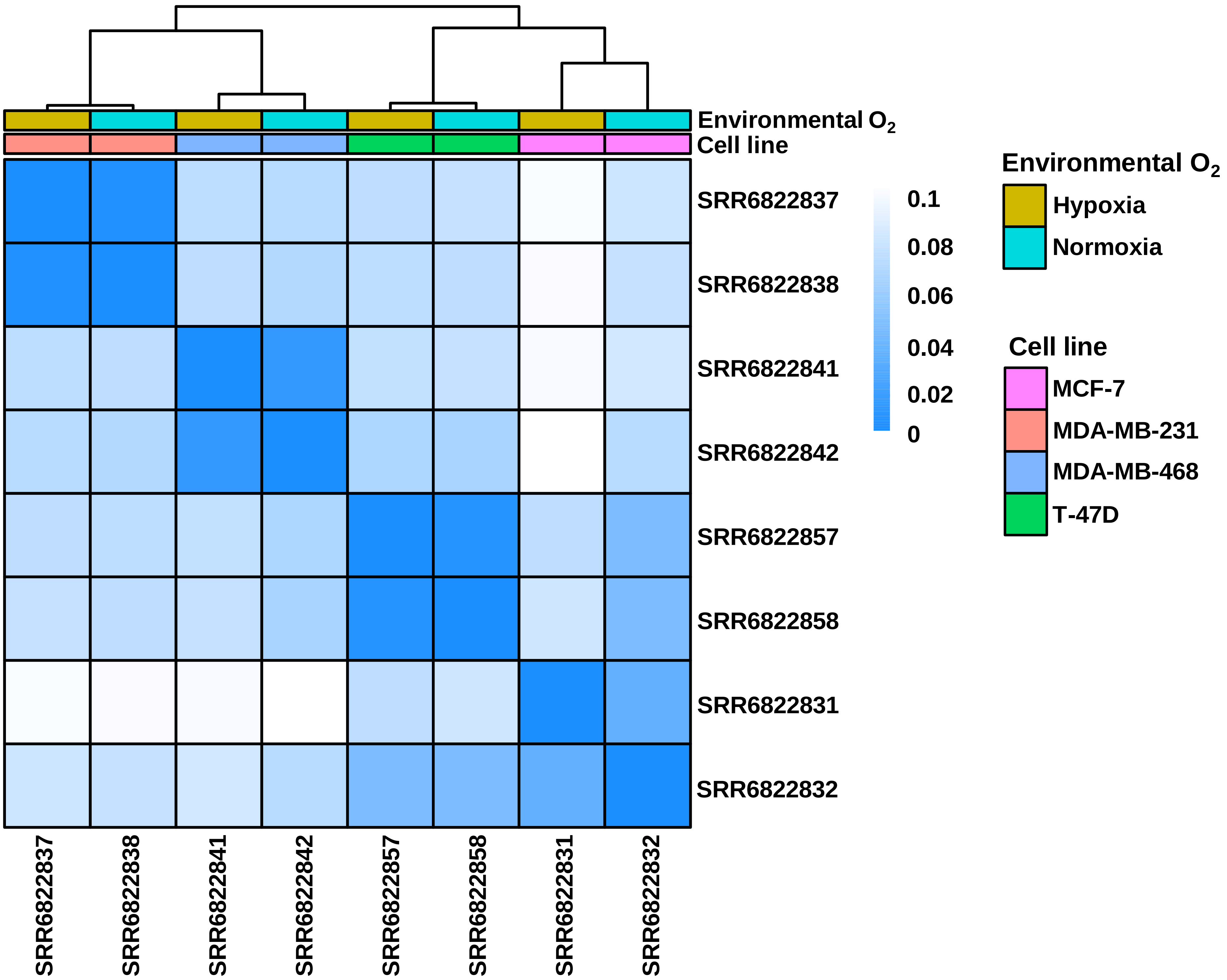


Supplementary Figure S1. Euclidean distance matrix (EDM) of squared distances between GSE111653 ERα+ and TNBC, hypoxic and normoxic breast cancer transcriptome datasets.

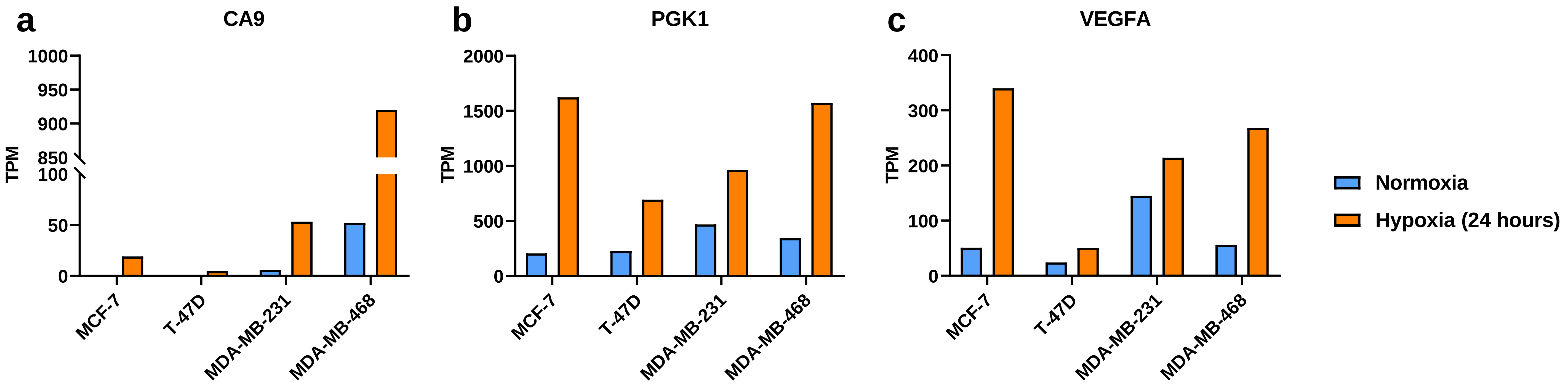


Supplementary Figure S2. High-throughput transcriptomic analysis of hypoxia-responsive genes (A) *CA9* and (B) *PGK1* and (C) *VEGFA* in breast cancer cell lines cultured in 20% $O_2$ (normoxia) or 1% $O_2$ (hypoxia) for 24 hours. The normalised read counts in transcripts per million (TPM) for each gene in MDA-MB-231, MDA-MB-468, MCF-7 and T-47D breast cancer cell lines cultured in normoxia (blue) or hypoxia (orange) are shown. Only one replicate was available for initial analysis.

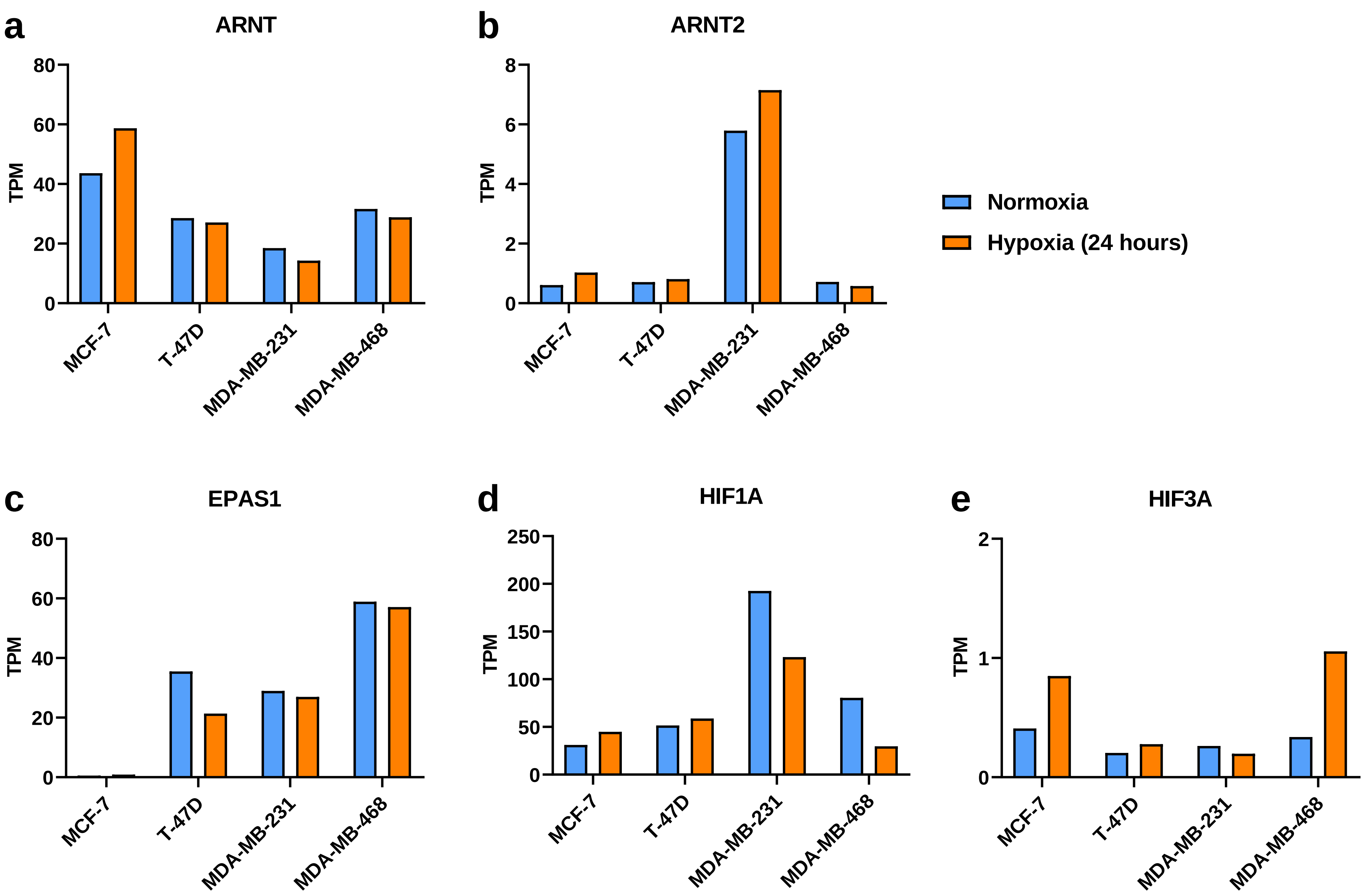


Supplementary Figure S3. High-throughput transcriptomic analysis of hypoxia-inducible factor (HIF) genes (A) *ARNT,* (B) *ARNT2,* (C) *EPAS1,* (D) *HIF1A* and (E) *HIF3A* in breast cancer cell lines cultured in 20% $O_2$ (normoxia) or 1% $O_2$ (hypoxia) for 24 hours. The normalised read counts in transcripts per million (TPM) for each gene in MDA-MB-231, MDA-MB-468, MCF-7 and T-47D breast cancer cell lines cultured in normoxia (blue) or hypoxia (orange) are shown. Only one replicate was available for initial analysis.

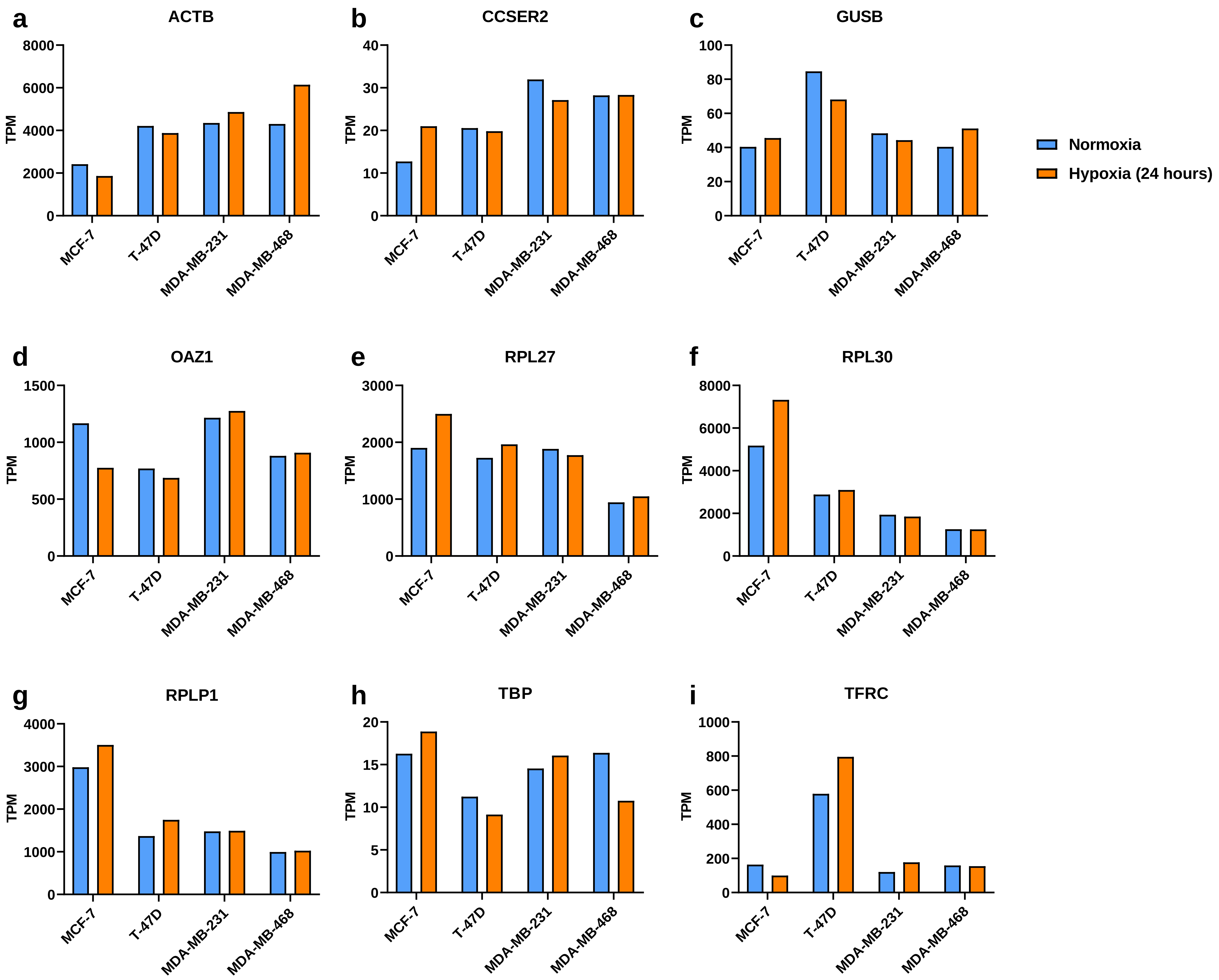

a
ACTB
b
CCSER2
c
GUSB
d
OAZ1
e
RPL27
f
RPL30
g
RPLP1
h
TBP
i
TFRC
TPM
MCF-7
T-47D
MDA-MB-231
MDA-MB-468
Normoxia
Hypoxia (24 hours)

Supplementary Figure S4. High-throughput RNA-seq analysis of RGs in breast cancer cell lines cultured in 20% $O_2$ (normoxia) or 1% $O_2$ (hypoxia) for 24 hours. A shortlist of RGs to be assessed for their suitability for RT-qPCR normalisation included (A) *ACTB*, (B) *CCSER2*, (C) *GUSB*, (D) *OAZ1*, (E) *RPL27*, (F) *RPL30*, (G) *RPLP1* (H) *TBP* and (I) *TFRC*. The normalised read counts in transcripts per million (TPM) for each gene in MDA-MB-231, MDA-MB-468, MCF-7 and T-47D breast cancer cell lines cultured in normoxia (blue), or hypoxia (red) are shown. Only one biological replicate was available for analysis.

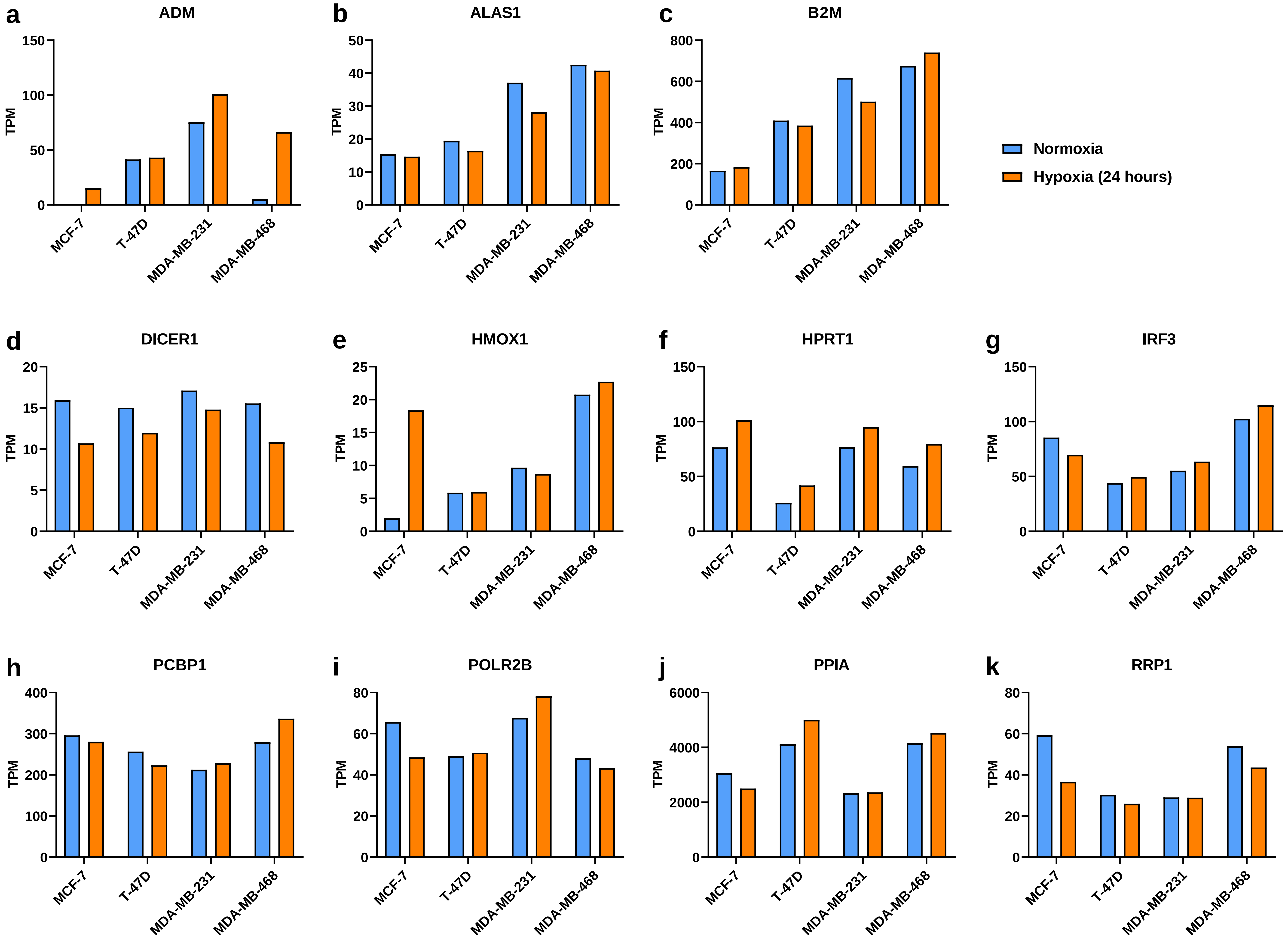

a
ADM
b
ALAS1
c
B2M
d
DICER1
e
HMOX1
f
HPRT1
g
IRF3
h
PCBP1
i
POLR2B
j
PPIA
k
RRP1
TPM
MCF-7
T-47D
MDA-MB-231
MDA-MB-468
Normoxia
Hypoxia (24 hours)

Supplementary Figure S5. High-throughput transcriptomic analysis of commonly used reference genes (A) *ADM,* (B) *ALAS1,* (C) *B2M,* (D) *DICER1,* (E) *HMOX1,* (F) *HPRT1,* (G) *IRF3,* (H) *PCBP1,* (I) *POLR2B,* (J) *PPIA* and (K) *RRP1* in breast cancer cell lines cultured in 20% $O_2$ (normoxia) or 1% $O_2$ (hypoxia) for 24 hours. The normalised read counts in transcripts per million (TPM) for each gene in MDA-MB-231, MDA-MB-468, MCF-7 and T-47D breast cancer cell lines cultured in normoxia (blue) or hypoxia (orange) are shown. Only one replicate was available for initial analysis.

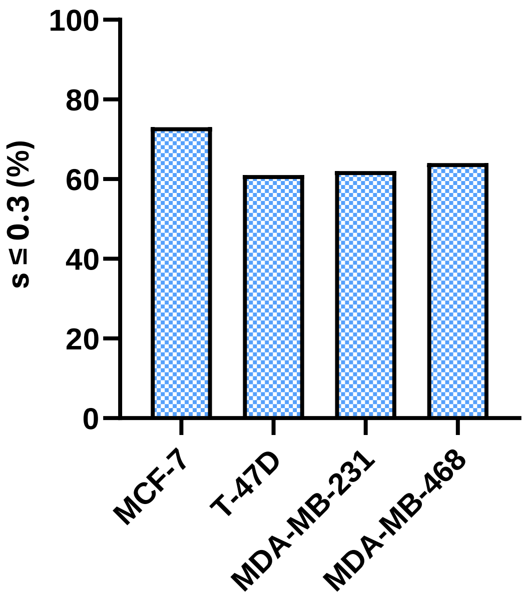


Supplementary Figure S6. *s* score analysis across all genes in breast cancer cell lines. For 30,187 genes, the percentage of genes that have an *s* score of equal to or greater than (≥) 0.3 has been calculated in each breast cancer cell line.

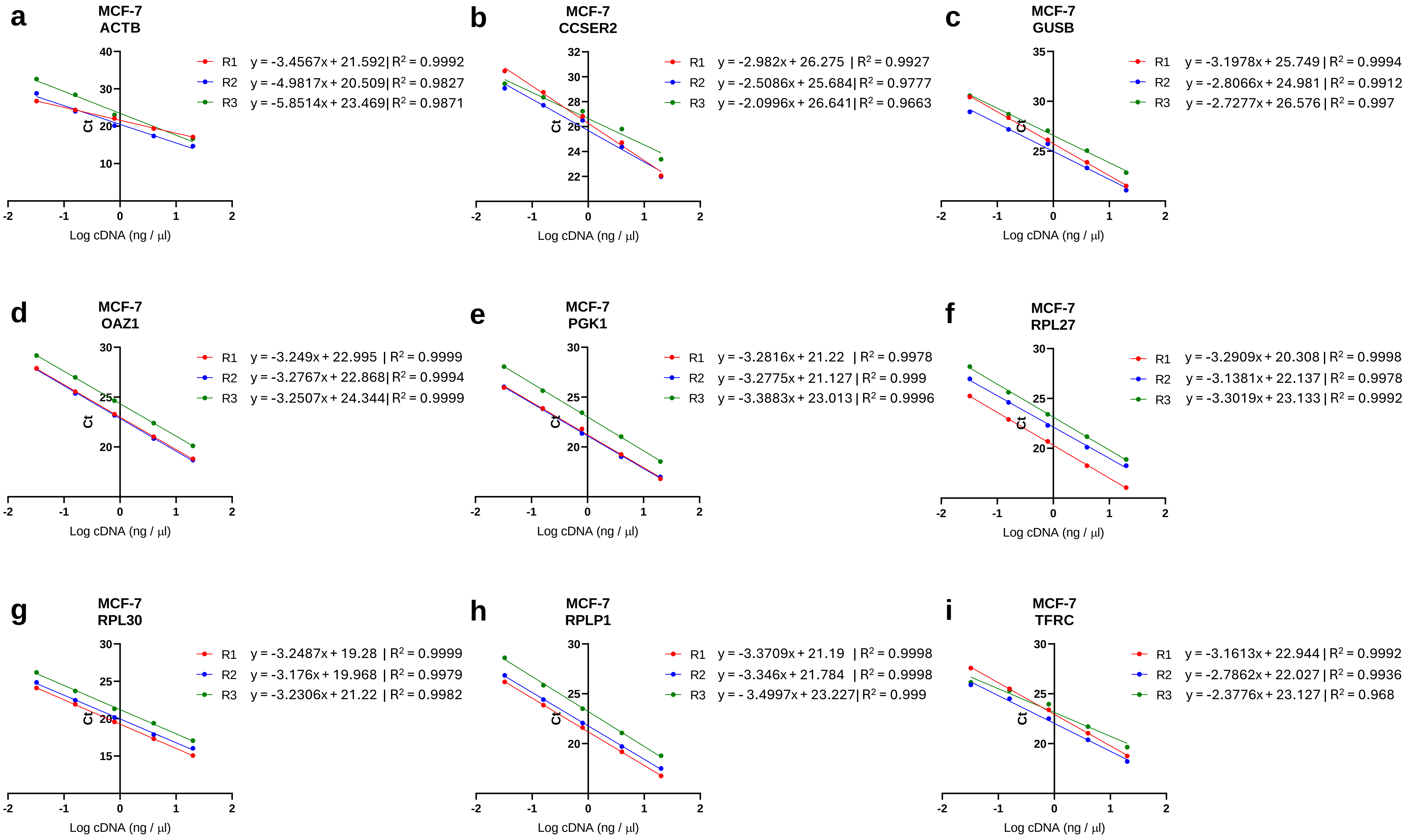


Supplementary Figure S7. Standard curves with line equations and $R^2$ value for (A) *ACTB,* (B) *CCSER2,* (C) *GUSB,* (D) *OAZ1,* (E) *PGK1,* (F) *RPL27,* (G) *RPL30,* (H) *RPLP1* and (I) *TFRC* in MCF-7 breast cancer cells. R1, R2 and R3 refers to the biological replicate.

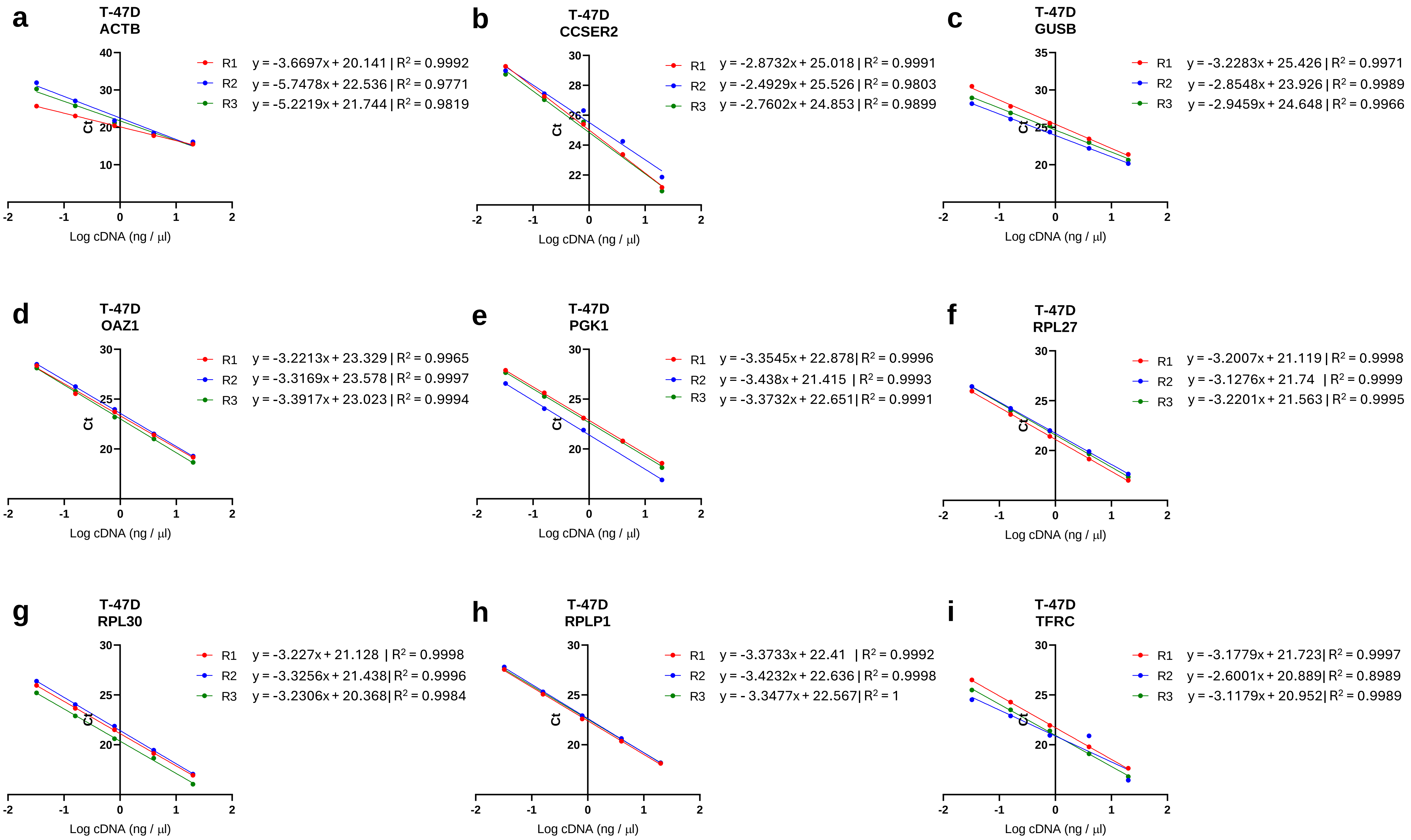


Supplementary Figure S8. Standard curves with line equations and $R^2$ value for (A) *ACTB,* (B) *CCSER2,* (C) *GUSB,* (D) *OAZ1,* (E) *PGK1,* (F) *RPL27,* (G) *RPL30,* (H) *RPLP1* and (I) *TFRC* in T-47D breast cancer cells. R1, R2 and R3 refers to the biological replicate.

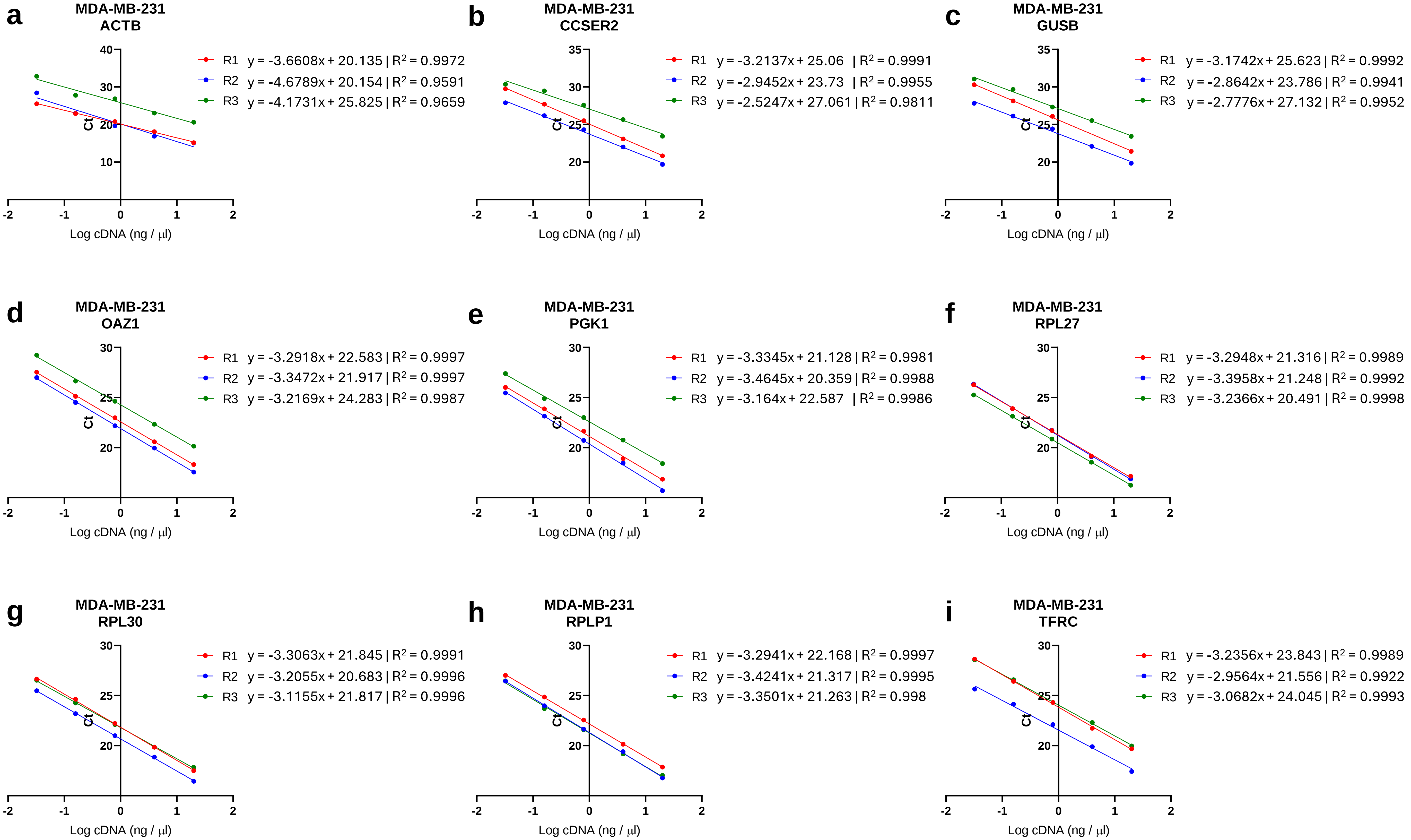


Supplementary Figure S9. Standard curves with line equations and $R^2$ value for (A) *ACTB,* (B) *CCSER2,* (C) *GUSB,* (D) *OAZ1,* (E) *PGK1,* (F) *RPL27,* (G) *RPL30,* (H) *RPLP1* and (I) *TFRC* in MDA-MB-231 breast cancer cells. R1, R2 and R3 refers to the biological replicate.

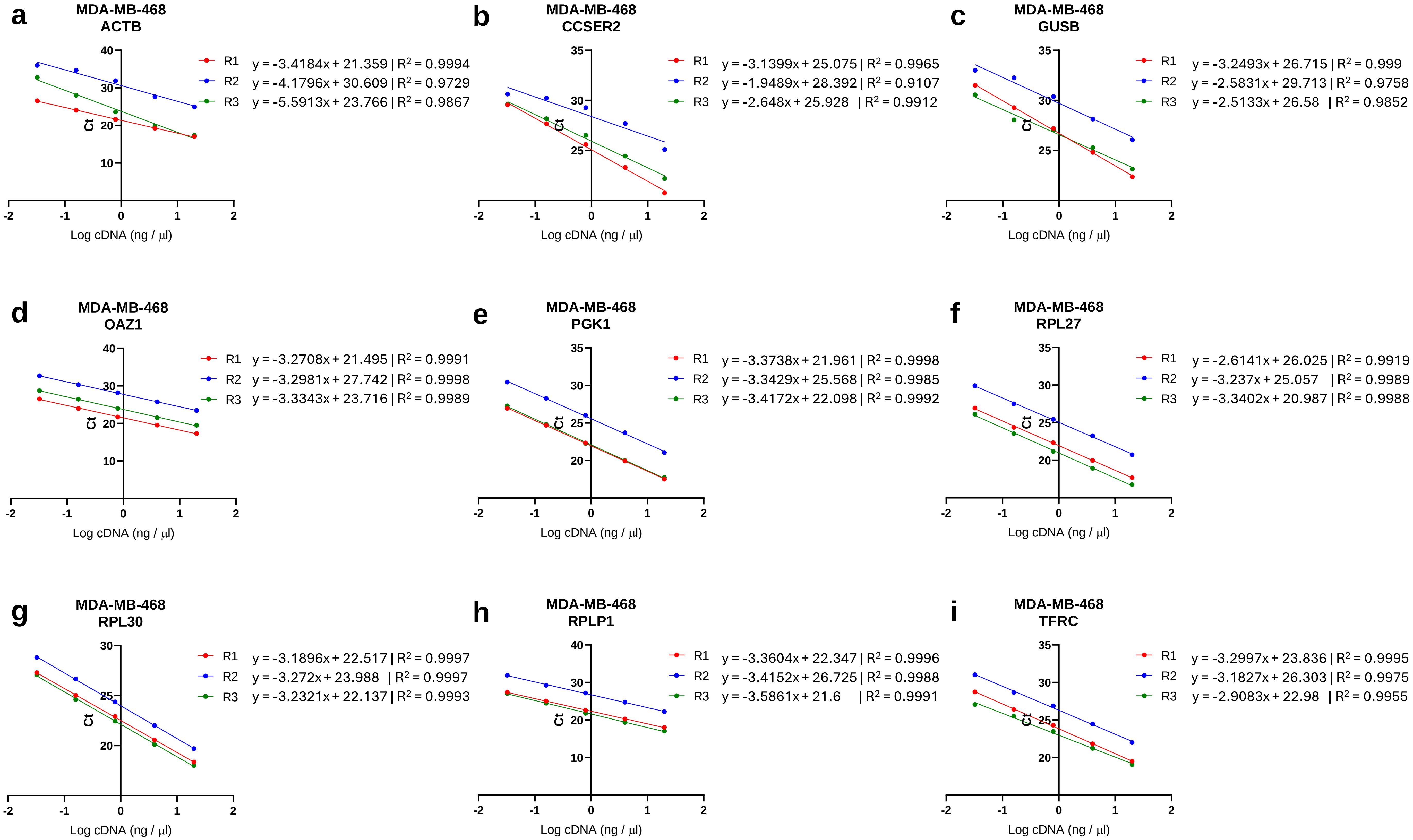


Supplementary Figure S10. Standard curves with line equations and $R^2$ value for (A) *ACTB,* (B) *CCSER2,* (C) *GUSB,* (D) *OAZ1,* (E) *PGK1,* (F) *RPL27,* (G) *RPL30,* (H) *RPLP1* and (I) *TFRC* in MDA-MB-468 breast cancer cells. R1, R2 and R3 refers to the biological replicate.

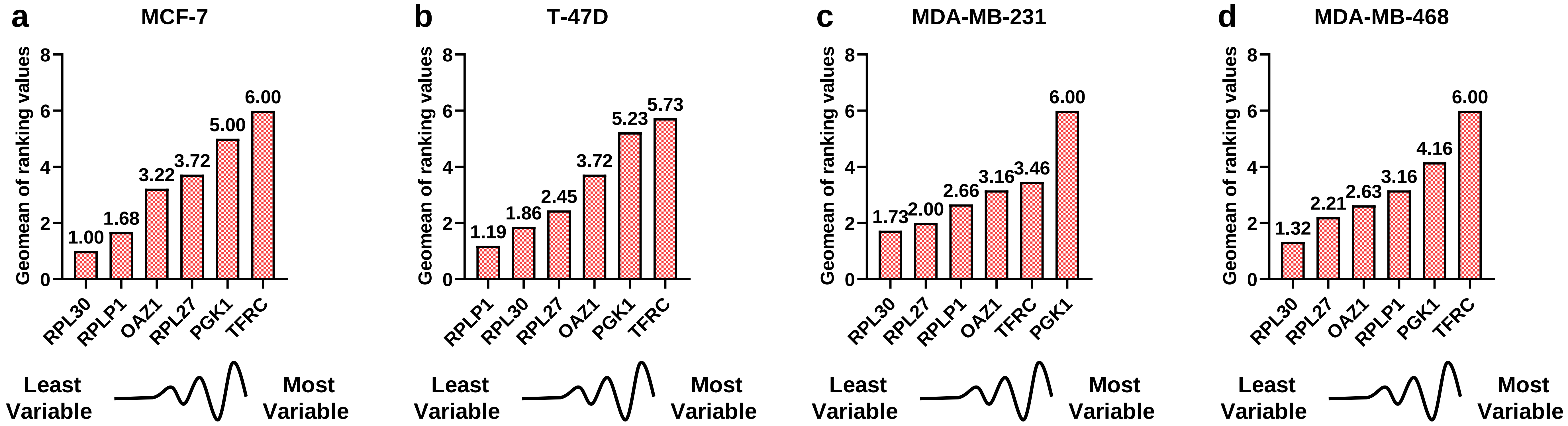


Supplementary Figure S11. Geometric mean (Geomean) of ranking values for each RG candidate according to RefFinder. The final overall ranking of RG candidates was determined by RefFinder based on the geometric mean of the weights of each gene from GeNorm, NormFinder, BestKeeper and the comparative ΔCt method for (A) MCF-7 (B) T-47D (C) MDA-MB-231 and (D) MDA-MB-468 breast cancer cell lines.
